# Diploid genome assembly of the Malbec grapevine cultivar enables haplotype-aware analysis of transcriptomic differences underlying clonal phenotypic variation

**DOI:** 10.1101/2023.11.30.569420

**Authors:** Luciano Calderón, Pablo Carbonell-Bejerano, Claudio Muñoz, Laura Bree, Cristobal Sola, Daniel Bergamin, Walter Tulle, Sebastian Gomez-Talquenca, Christa Lanz, Carolina Royo, Javier Ibáñez, José Miguel Martinez-Zapater, Detlef Weigel, Diego Lijavetzky

**Affiliations:** Instituto de Biología Agrícola de Mendoza (CONICET-UNCuyo), Genetica y Genomica de Vid. (5505) Mendoza, Chacras de Coria, Argentina; Instituto de Ciencias de la Vid y del Vino, ICVV, CSIC - Universidad de La Rioja - Gobierno de La Rioja. (26007) La Rioja, Logroño, Spain; Department of Molecular Biology, Max Planck Institute for Biology Tübingen. (72076) Tübingen, Germany; Facultad de Ciencias Agrarias (UNCuyo), Cátedra Fitopatología. (5505) Mendoza, Chacras de Coria, Argentina; Vivero Mercier Argentina. (5500) Mendoza, Perdriel, Argentina; Instituto Nacional de Tecnología Agropecuaria, Plant Virology Laboratory. (5534) Mendoza, Luján de Cuyo, Argentina

## Abstract

To preserve their varietal attributes, established grapevine cultivars (*Vitis vinifera* L. ssp. *vinifera*) must be clonally propagated, due to their highly heterozygous genomes. Malbec is a France-originated cultivar appreciated for producing high-quality wines, and is the offspring of cultivars Prunelard and Magdeleine Noire des Charentes. Here, we have built a diploid genome assembly of Malbec, after trio binning of PacBio long reads into the two haploid complements inherited from either parent. After haplotype-aware deduplication and corrections, complete assemblies for the two haplophases were obtained with very low haplotype switch-error rate (<0.025). The haplophases alignment identified >25% of polymorphic regions. Gene annotation including RNA-seq transcriptome assembly and *ab initio* prediction evidence resulted in similar gene model numbers for both haplophases. The annotated diploid assembly was exploited in the transcriptomic comparison of four clonal accessions of Malbec that exhibited variation in berry composition traits. Analysis of the ripening pericarp transcriptome using either haplophases as reference yielded similar results, although some differences were observed. Particularly, among the differentially expressed genes identified only with the Magdeleine-inherited haplotype as reference, we observed an over-representation of hypothetically hemizygous genes. The higher berry anthocyanin content of clonal accession 595 was associated with increased abscisic acid responses, possibly leading to the observed overexpression of phenylpropanoid metabolism genes and deregulation of genes associated to abiotic stress response. Overall, the results highlight the importance of producing diploid assemblies to fully represent the genomic diversity of highly heterozygous woody crop cultivars and to unveil the molecular bases of clonal phenotypic variation.

## 2. Introduction

The cultivated grapevine (*Vitis vinifera* L. subsp. *vinifera*) was one of the first plant species accounting for a reference genome ^1^. To facilitate genome assembly, this reference was produced from a nearly homozygous line (i.e. PN40024) obtained after several rounds of selfing of the cultivar Helfensteiner, the result of Pinot noir and Schiava grossa outcross ^1,2^. The PN40024 reference genome has been a key tool for the study of grapevine molecular biology and genetics and both the reference assembly and annotation have been continuously improved ^2–4^.

The genomes of grapevine cultivars are characterized by enormous diversity, including the many differences that typically distinguish the two haplotypes of each cultivar ^7^. On average one single nucleotide polymorphism (SNP) every 100 bp and one Insertion-deletion (InDel) difference every 450 bp ^7,8^. Also, larger structural variants (SVs) ≥50 bp have been reported to affect ∼17% of the genome space of each haplotype within the same cultivar ^9^. Hemizygosity is also rampant in grapevines, since many SVs cause presence-absence variation (PAV) between haplotypes ^8^, which may have severe phenotypic impacts ^10^. In addition to the PN40024 reference genomes, several additional cultivars have seen their genomes assembled from Pacific Biosciences (PacBio) long reads using the haplotype-aware assemblers FALCON-unzip ^6,9,11–14^. FALCON-unzip outputs pseudo-haplotypes, which usually represent a single allele at each position without preserving phase information across the genome ^15^, although tools such as HaploSync have been developed to completely phase this type of assemblies ^16^. In this context, the combination of multiple sequencing technologies such as Oxford Nanopore Technologies (ONT) and short reads^21^, the use of PacBio HiFi and newer algorithms such as HiFiasm ^17^, as well as information from parent-offspring trios ^15^ have notably improved assembly phasing and quality of grapevine genomes ^4,18 19^.

The high and uneven heterozygosity of cultivated grapevine genomes is most likely the consequence of the outcrossing nature of their dioecious wild ancestor (*V. vinifera* ssp. *sylvestris*) and their vegetative way of propagation ^6^. Moreover, cultivars relevant for wine making have originated from complex processes including the outcrossing of preexisting genetically divergent cultivars ^20^ and eventual post-domestication hybridization with local wild relatives of the subspecies *sylvestris* ^21,22^. Irrespective of their origin, progenies with desired phenotypic traits have been vegetatively propagated to retain those attributes. Although vegetative propagation mostly ensures cultivar uniformity, it also favors the accumulation of somatic mutations in the absence of outcrossing ^23^. In fact, somatic variation is known to exist at the phenotypic ^24^, genetic ^13^ and transcriptomic ^25^ levels. Moreover, within each grape cultivar growers select somatic variants to propagate them, labelling these selections as clones. Therefore, somatic diversity has been relevant for viticulture as a source of trait innovation and adaptation. Because the global wine market is very slow at accepting new outcross varieties, a deep understanding of the variation that exists within elite varieties is crucial for intra-cultivar improvement and to meet emerging challenges from climate change ^26^.

Malbec, a red cultivar appreciated for its potential to produce high-quality wines, originated in the Cahors region of France (where it is known as Cot) as the result of an outcross of cultivars Magdeleine Noir des Charentes (hereafter Magdeleine) and Prunelard ^27^. Malbec was introduced to Argentina in the 1850s, becoming the most relevant cultivar for the wine industry of this country ^28^. Currently, Argentina harbors the largest planted surface of Malbec in the world ^29^ and 85% of this production is concentrated in Mendoza province ^30^. Since its introduction, this cultivar has shown remarkable adaptability, producing wines with properties that reflect the different growing conditions ^31^. The observed adaptability may be linked to the phenotypic plasticity exhibited by Malbec, as clonal variability has been reported for traits relevant for the industry, such as polyphenolic composition of berries ^32^ and phenology ^33^. Also molecular plasticity has been described among clones, since epigenetic ^34^ and genetic ^35^ variability has been reported.

The main objective of this study was to produce a phased and annotated genome assembly of Malbec representing both haploid complements, or haplophases, to serve as reference for intra-cultivar variation studies. We used our assembly in a transcriptome analysis to investigate the molecular bases of Malbec clonal variants that differ in fruit traits relevant for wine making. The transcriptomic analysis was performed following a haplotype-aware scheme and the results obtained using separately each assembled haplophase as reference were compared. This allowed us to further evaluate potential reference biases, and to identify haplotype-specific differentially expressed genes that could be correlated with clonal variation in berry composition.

## 3. Results

### The diploid *de novo* genome assembly of Malbec

A total of 82.3 Gb of sequence coverage (168x coverage of the haploid genome) was produced for Malbec in 4,117,439 PacBio CLR reads, with a mean read length of 20 kb and read N50 of 33.7 kb (Table S1A). Illumina short reads were obtained for Malbec (13.4 Gb) as well as the parental cultivars Prunelard (46.9 Gb) and Magdeleine (47.6 Gb) (Table S1B). Malbec PacBio reads were binned into two sets of 35.9 Gb and 46.0 Gb, corresponding to the haplotypes inherited from Magdeleine and Prunelard (Table S2A). The binned PacBio long reads were separately assembled into two haplophases, named hereafter Malbec-Pru for the inherited from Prunelard and Malbec-Mag for the inherited from Magdeleine. The draft raw assemblies of Malbec-Pru and Malbec-Mag had good contiguity, with contig N50 = 6.2 Mb and 5.6 Mb, and gene completeness of 98.8% and 98.2% of BUSCO core genes represented (Table S2B). However, the draft assemblies were larger than the expected for a haploid grapevine genome of ∼475 Mb ^2^, namely 608 Mb for Malbec-Pru and 558 Mb for Malbec-Mag, and they had a high percentage of duplicated BUSCO genes, 21.7% and 10.4% (Table S2B). After haplotype-aware deduplication, consensus polishing and miss-assembly corrections, the two haplophases were closer to the expected size, 479.3 Mb and 479 Mb, with improved contiguity of N50 = 7.7 Mb and 6.6 Mb. Additionally, we obtained a smaller percentage of duplicated BUSCO genes (i.e., 2.7% and 1.9%) without a significant change in completeness: 98.6% and 98.1% (Table 1A). The polished and deduplicated haplophases were reference-based scaffolded, resulting in 19 pseudomolecules, as expected for the haploid set of grapevine nuclear chromosomes (Fig. S1), with a scaffold N50 = 24.4 for Malbec-Pru and 24.5 Mb for Malbec-Mag (Table 1B). Some unplaced contigs remained after the scaffolding, for Malbec-Pru = 21 contigs (total length = 3.3 Mb) and for Malbec-Mag = 32 contigs (total length = 5.74 Mb). Malbec-Mag corresponds to the maternally inherited haplotype ^27^, and included one scaffold for a partial mitochondrial chromosome (length = 115,997 bp). Finally, a group of contigs of short length (N50 = 0.2 Mb), that were slightly over-represented in repetitive sequences and under-represented in BUSCO complete genes (avg. 17%) compared to the assembly size, were discarded from the final assembly after the deduplication process (Table S2C).

**Table 1.**
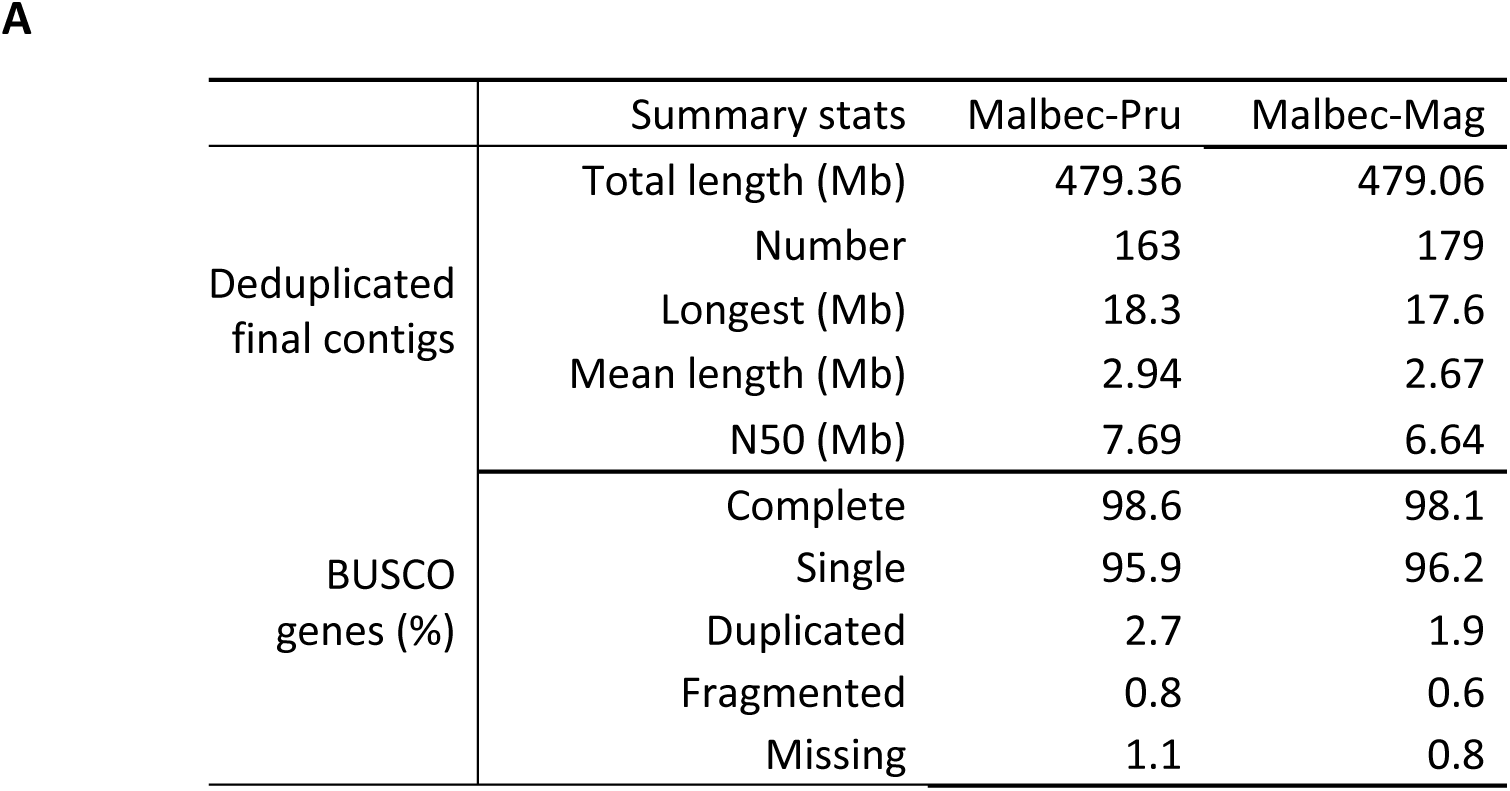
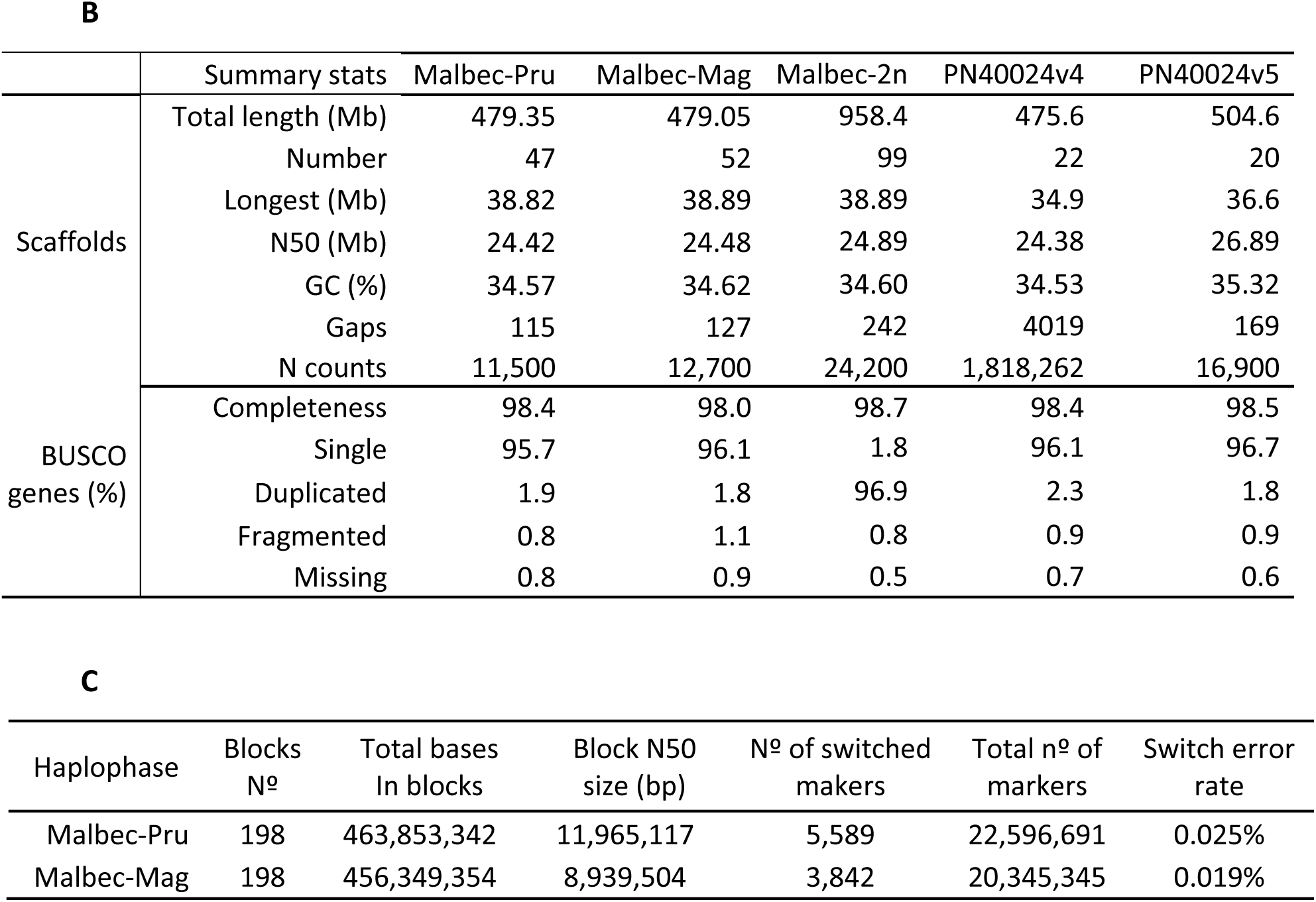
Summary statistics obtained for the Malbec diploid genome assembly after deduplication pipeline. (A) Results obtained for Malbec-Pru and Malbec-Mag contigs. (B) Results obtained after scaffolding process for both Malbec haplophases and the diploid (2n) assembly, compared to the PN40024.v4 (REF) and PN40024.v5 (T2T) reference genome assembly versions. Percentages are expressed over a total of 2,326 BUSCO genes in the eudicots lineage. (C) Evaluation of the phasing process performed by comparing *k-mers* between the Malbec haplophase assemblies and raw short-reads from the parental cultivars.

Phasing accuracy and consensus quality was assessed on the final scaffolded assemblies. The Malbec-Pru assembly comprised 97.3% of the Prunelard-specific *hap-mers* and only 0.13% of Magdeleine-specific ones (Fig. 1A), whereas the Malbec-Mag assembly comprised 98.1% of the Magdeleine-specific *hap-mers* and only 0.03% of the Prunelard-specific ones (Fig. 1B). The consensus quality score (QV) was estimated for Malbec-Pru = 41.62 (error rate = 6.9 e-5) and Malbec-Mag = 42.06 (error rate = 6.2 e-5) (Table 1C). Total bases placed in proper phase blocks and the phase blocks N50 for each scaffolded haplophase assembly were for Malbec-Pru = 463.8 Mb (N50 = 11.96 Mb) and Malbec-Mag = 456.3 Mb (N50 = 8.9 Mb) (Figs 1A and B, Fig. S2). Also, a very low haplophase switch error rate was observed, 0.025% for Malbec-Pru and 0.019% for Malbec-Mag (Table 1C).

**Figure 1.**
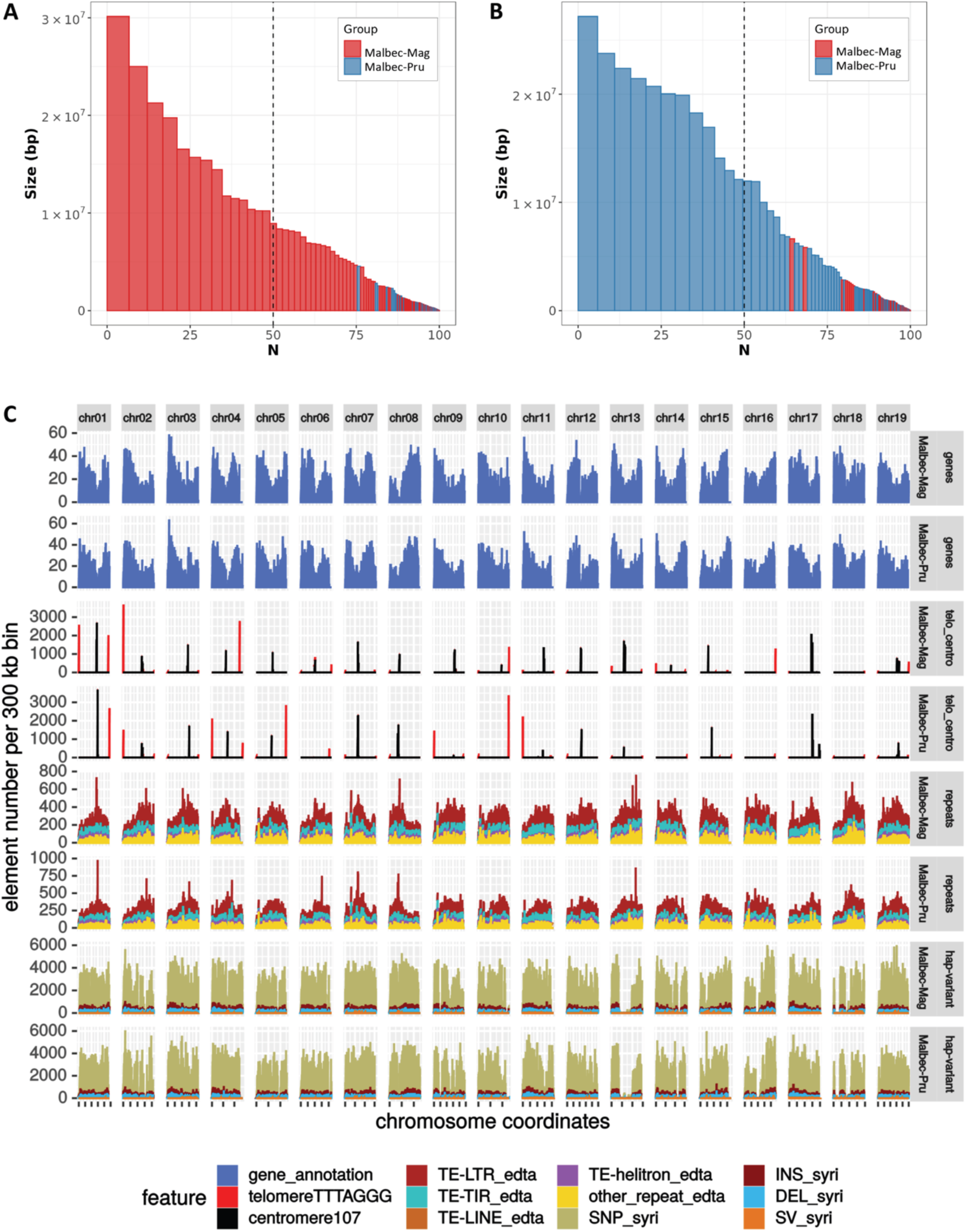
Barplots of the phased blocks sorted by size to visualize the efficiency of the phasing process for (A) Malbec-Mag and (B) Malbec-Pru haplophases. For the former the red bars are the own blocks and the blue bars are the switched blocks, for the latter the color code is inverted. Vertical bars represent phase blocks, defined as a continuous set of *k-mers* inherited from the same haplotype (i.e., *hap-mers*). The *X*-axis is the percentage of the genome size covered by blocks of this size of larger, expressed in base pairs (*Y*-axis). (B) Distribution of the annotated features along the 19 scaffolded pseudomolecules for Malbec haplophases, including: gene density, centromeres, telomeres, transposable elements (LTRs, TIR, LINE and Helitron) and haplotype variants (SNPs, INDELs, SVs).

The estimated divergence between the two haplophases did not change considerably based on which one was used as reference, and we report the results obtained with Malbec-Pru as reference and Malbec-Mag as query. Sequence differences affected 7.09% (33.55 Mb) of Malbec-Mag, this included SNPs, InDels, copy number variations (CNVs), highly divergent regions (HDRs) and tandem repeats (TRs) (Fig. 1C and Table S3). Considering the accumulative length, HDR was the most abundant type of sequence variation and duplications (DUPS) were the most abundant type structural variation, affecting 4.9% (23.28 Mb) and 3.98% (18.84 Mbp) of Malbec-Mag (Fig. 2A). SNPs and InDels occurred every 147 and 1458 nt, respectively; and SNPs distribution along the chromosomes was rather uneven (Fig. 1C). Several drops of SNP density were detected, possibly as consequence of runs of homozygosity. The longest drop of SNPs occurred on chr13 (Fig. 1C) and this overlapped with a large inversion between the haplophases (Fig. 2A), which may appear as a SNP desert in the SyRI output ^36^. Structural variations affected 9.97% (47.2 Mb) of Malbec-Mag, more precisely inversions (INV) = 2.07%, translocations (TRANS) = 3.92% and duplications (DUPS) = 3.98% (Fig. 2A and Table S3A). Finally, 76.5% (362.14 Mb) of the haplophases were identified as syntenic and 14.2% (67.2 Mb) were classified as not aligned (Fig. 2A and Table S3A), altogether indicating for a high heterozygosity of Malbec cultivar represented in the diploid assembly. Moreover, in both haplophases an average of 29% of genes were affected by at least one SV (namely: TRANS, INV, DUP, NOTAL) (Table S3C). Furthermore, genes overlapping with SVs were over-represented in biological processes GO terms associated to ‘defense response’, ‘external biotic stimulus’ and ‘DNA metabolic processes’ in both haplophases. At the same time, ‘valine biosynthetic process’ and ‘phosphorylation’ were enriched solely in Malbec-Pru and ‘glucan biosynthetic process’ was enriched in Malbec-Mag (Table S3D).

**Figure 2.**
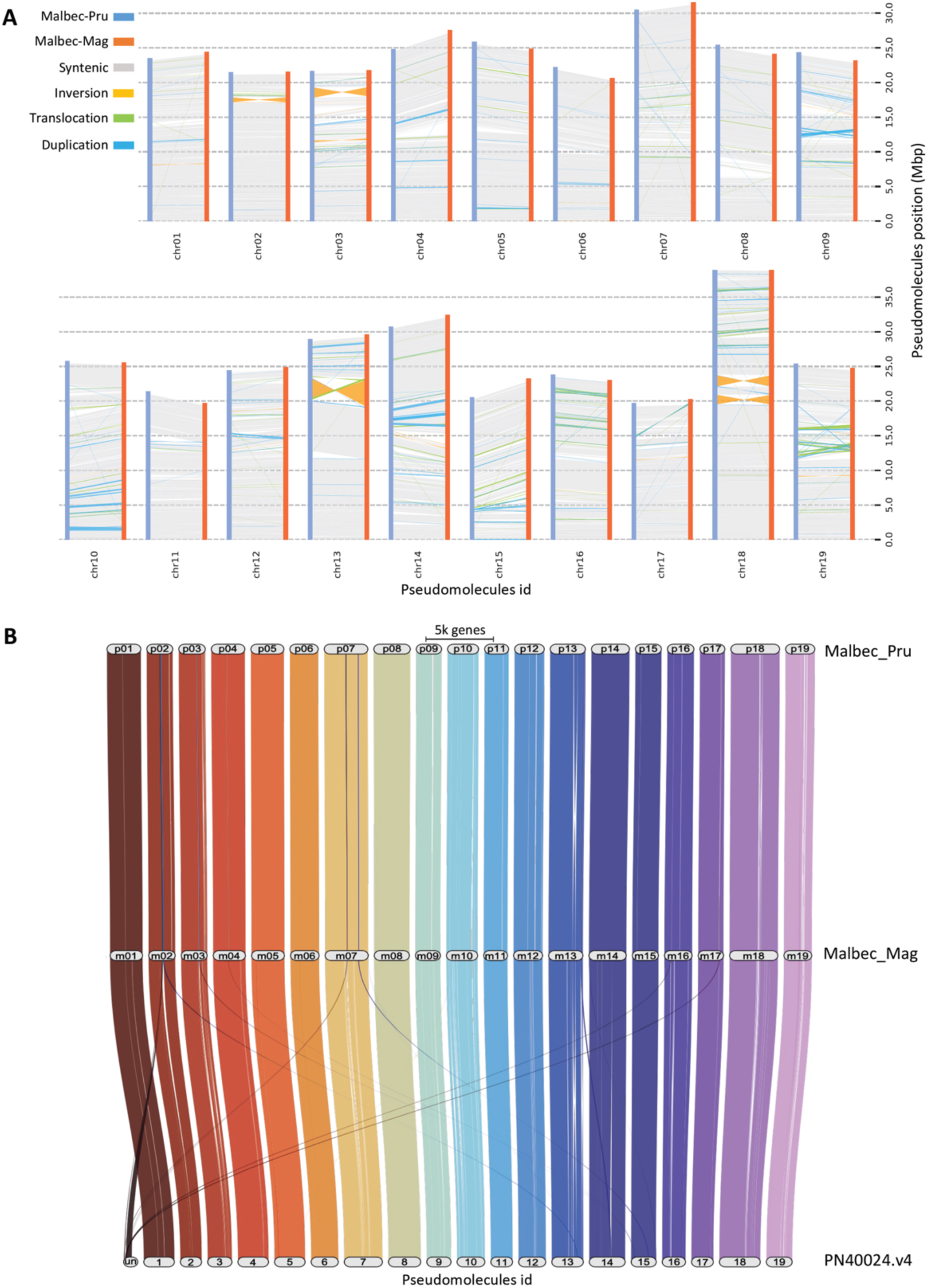
Synteny analysis based on the comparison of the 19 pseudomolecules assembled for Malbec haplophases. (A) Structural variations (SVs) distribution along pseudomolecules, with Malbec-Pru (blue) considered as the reference and Malbec-Mag (orange) as the query. Results are based on the percentages estimated for the query, 76.5% (362.14 Mb) of the assembly resulted syntenic and 9.97% (47.2 Mb) was affected by different types of SVs (inversions, translocations and duplications). (B) Riparian plots representing the synteny of orthologous genes along the 19 scaffolded pseudomolecules for each haploid complement of Malbec. A strong synteny was observed among the assembled pseudomolecules of Malbec haplophases and the grapevine reference genome (PN40024.v4). Also, genes contained in the “Unknown” (Un) chromosome of the PN40024.v4 exhibited orthologous genes assigned to different pseudomolecules in Malbec haplophases.

### Gene model predictions for the assembled haplophases

Annotation and soft-masking of transposable elements (TEs) was performed on the Malbec diploid assembly, after concatenating the two haplophases (958 Mb). Almost half, 48.4 % (464 Mb), of the genome accounted for TEs, including structurally intact and fragmented TEs (Table S4). The great majority was classified as long terminal repeats (LTRs) representing 28.8% (276 Mb) of the genome, whereas Gypsy was the most abundant type among the LTRs class with 13.36% (128 Mb) (Fig. 1C, Table S4). In addition, telomeric and centromeric repeats were identified in the diploid assembly. Out of the 19 scaffolded pseudomolecules for each haplophase, 14 and 18 featured telomeric repeats in at least one end in Malbec-Mag and Malbec-Pru (Fig. 1C). Centromeric repeats were also detected (between 173 and 5817 clustered copies) in most pseudomolecules, with the exception of chr16 and chr18 for both haplophases, and for chr06, chr10 and chr14 for Malbec-Pru. The fact that the annotated position of centromeric repeats overlapped between the two haplophases, as well as with LTR-TEs, suggest that these regions might correspond to the -at least partially-assembled centromeres (Fig. 1C).

The final weighed consensus gene models obtained for the Malbec diploid genome assembly yielded similar metrics to those of the highly curated PN40024.v4 ^2^ annotations, whereas the annotations of the T2T PN40024.v5 assembly ^4^ differed in parameters like exon and UTR length (Table 2). The number of protein coding loci was 35,739 for Malbec-Pru and 35,463 for Malbec-Mag, with an average of 1.2 transcript isoforms per locus (Table 2). Annotated genes were similarly distributed when comparing the pseudomolecules of both haplophases. Drops of gene density overlapped with centromeric repeats, and the former were also evident even in those pseudomolecules where peaks of centromeric repeats were not detected, like chr16 and chr18 (Fig. 1C), further supporting the putative position of the centromeres.

**Table 2.** Coding protein sequences predicted for Malbec haplophases (considering isoforms), compared to the PN40024.v4 (REF) and the PN40024.v5 (T2T) annotations.

|  | Malbec-Pru |  | Malbec-Mag |  | PN40024.v4 (REF) |  | PN40024.v5 (T2T) |  |
| --- | --- | --- | --- | --- | --- | --- | --- | --- |
|  | Total number | Mean length (bp) | Total number | Mean length (bp) | Total number | Mean length (bp) | Total number | Mean length (bp) |
| Protein coding genes | 35,739 | 5,086 | 35,463 | 5,071 | 35,197 | 4,742 | 37,476 | 6,133 |
| Transcripts | 44,590 | 5,025 | 44,183 | 5,017 | 41,160 | 4,689 | 40,994 | 6,494 |
| Exons | 236,022 | 319 | 233,585 | 320 | 208,719 | 283 | 179,096 | 645 |
| CDS | 44,590 | 1,182 | 44,183 | 1,180 | 41,160 | 1,126 | 40,994 | 1,235 |
| 5' UTRs | 21,386 | 371 | 21,221 | 372 | 17,478 | 280 | 23,267 | 1,327 |
| 3' UTRs | 22,138 | 670 | 21,818 | 676 | 18,344 | 440 | 23,278 | 1,459 |

Considering all annotated isoforms, the total number of predicted proteins for Malbec haplophases were higher (Malbec-Pru = 44,590 and Malbec-Mag = 44,183) than the one reported for the PN40024.v4 and v5 annotations (Table 3A). However, the size metrics such as the N50 of the predicted proteins were more similar (Table 3A). BUSCO completeness for the proteins annotated on each haplophase was close to 97%, whereas 98.6% completeness was achieved when all proteins annotated in the diploid assembly (Malbec-2n) were analyzed (Table 3B).

**Table 3.** (A) Summary metrics and (B) BUSCO completeness analysis of the proteins predicted in the genome assemblies. Results retrieved for each Malbec haplophase and for the diploid (2n) assembly are compared to the PN40024.v4 (REF) and PN40024.v5 (T2T) annotations that were also analyzed as quality control.

| Predicted proteins | Malbec-Pru | Malbec-Mag | Malbec-2n | PN40024 v4 (REF) | PN40024 v5 (T2T) |
| --- | --- | --- | --- | --- | --- |
| Total proteins | 44,590 | 44,183 | 88,773 | 41,160 | 40,994 |
| Min. length (aa) | 37 | 41 | 37 | 50 | 18 |
| Average length (aa) | 393 | 393 | 393 | 375 | 411 |
| Max. length (aa) | 5,480 | 5,488 | 5,488 | 5,481 | 66,180 |
| N50 (aa) | 532 | 534 | 533 | 519 | 543 |

**B**
| BUSCO proteins (%) | Malbec-Pru | Malbec-Mag | Malbec-2n | PN40024 v4 (REF) | PN40024 v5 (T2T) |
| --- | --- | --- | --- | --- | --- |
| Completeness | 96.8 | 96.7 | 98.6 | 98.7 | 95.8 |
| Single | 72.7 | 72.1 | 3.8 | 80.3 | 88.7 |
| Duplicated | 24.1 | 24.6 | 94.8 | 18.4 | 7.1 |
| Fragmented | 1.2 | 1.2 | 0.4 | 0.7 | 1.1 |
| Missing | 2.0 | 2.1 | 1.0 | 0.6 | 3.1 |

The orthology analysis based on predicted proteins showed, as expected ^2,27,37^, that there were more orthologues between Malbec haplophases than with the more complete PN40024 v4 reference genome predictions (Table S5A). Orthology was compared to the PN40024 v4 annotation, since it was more complete than the annotations available for the v5 (Table 3B). When only Malbec haplophases were compared ∼96% of the proteins could be placed in orthogroups, on average 4.1% of the proteins remained unassigned, and 2.45% were haplotype-specific orthogroups (Table S5B). The liftoff analysis, based on annotated genes, between both haplophases showed that there were 1,915 genes of the Malbec-Mag that were unmapped in Malbec-Pru and 2,296 unmapped genes for the inverse comparison. Interestingly, the GO enrichment analysis of unmapped genes suggested that the trihydroxystilbene synthase activity and the RNA (uridine-N3-)-methyltransferase molecular functions were overrepresented in Malbec-Mag and Malbec-Pru, respectively (Fig. S3). The riparian plots for synteny visualization, showed that most gene orthogroups matched in their physical location across the 19 pseudomolecules of Malbec haplophases and PN40024.v4 (Fig. 2B). Furthermore, many genes located in the Unknown chromosome of the PN40024.v4 had orthologues in genes assigned to chrMT, chr02, chr07, chr16 and chr17 in Malbec-Mag and chr02, chr03, chr07 and chr17 in Malbec-Pru (Fig. 2B and Fig. S4).

### Malbec haplophases retrieved congruent -although not identical- transcriptomic differences, that resembled the clonal phenotype variation

The obtained diploid assembly was employed to further understand the molecular bases of Malbec intra-cultivar phenotypic variation for berry composition traits. Phenotyping of 27 clones of Malbec grown in the same plot showed significant diversity for traits evaluated on mature berries. In a principal component analysis (PCA), the first component (PC1) explained 46.2% of the variation, mainly driven by total anthocyanins (TA), total polyphenols (TP) and must pH, whereas PC2 explained 27.3% of the variation and appeared to be associated with sugar content (Bx) and total acidity (Ac) (Fig. S5A). From the starting 27 phenotyped accessions, four clones (595, 596, 505 and 136N) were selected for further transcriptomic analysis (Fig. S5B). The greatest phenotypic differences among the selected clones were associated with total polyphenols (TP) and total anthocyanin (TA) concentrations, with accessions 595 and 596 having the highest and lowest values (*p*<0.05), whereas 505 and 136N had intermediate values (Fig. 3A). These differences among clones were consistent across seasons, despite the inter-annual differences in the TP and TA absolute values (higher in 2018 than in 2019) (Fig. 3B). Overall, accession 595 showed the highest TP and TA concentrations in mature berries at two consecutive harvest seasons.

**Figure 3.**
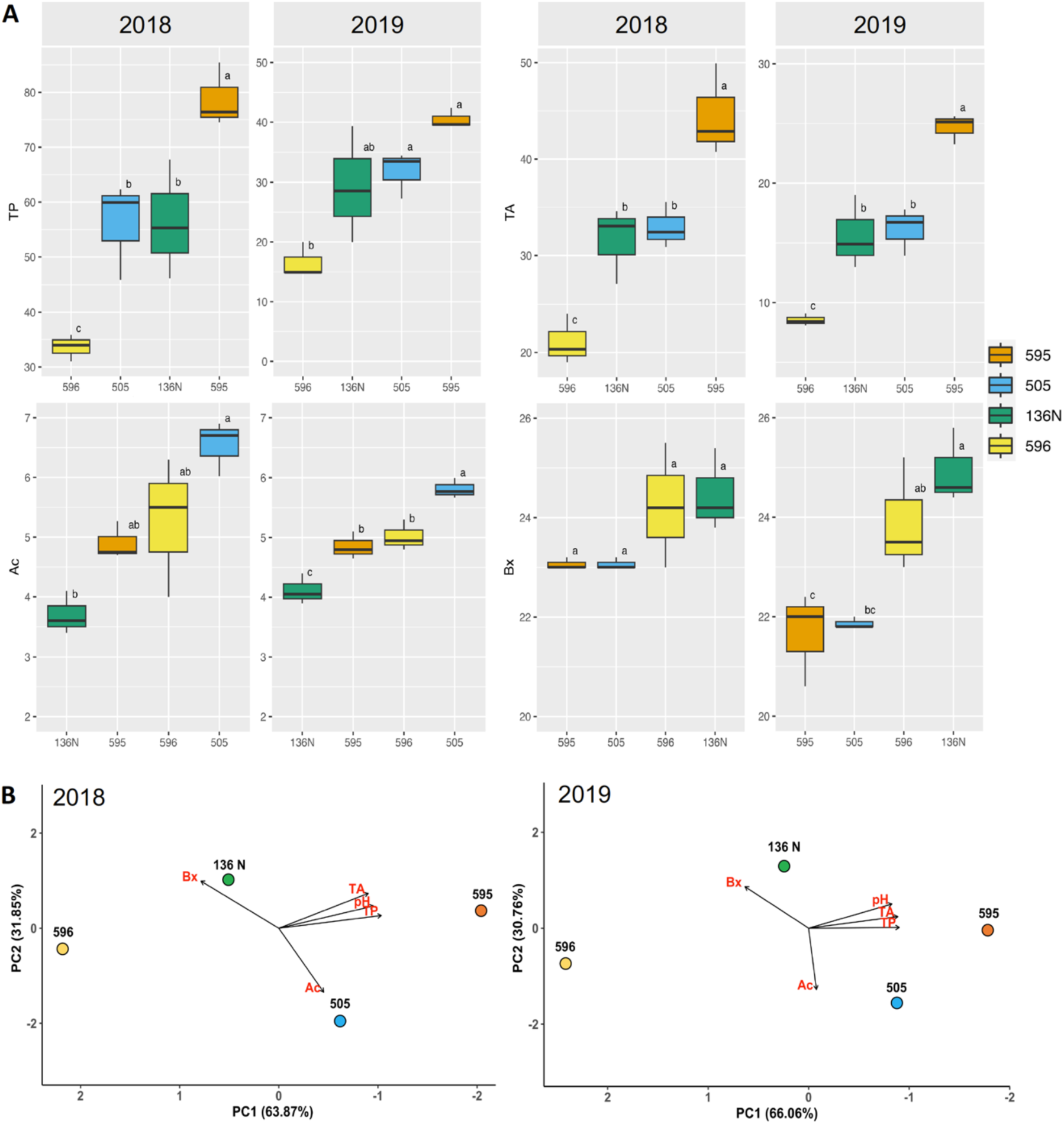
Phenotypic analyses conducted on the four clones selected to perform RNAseq experiments. All phenotypic data was obtained from mature berries during two consecutive years, 2018 and 2019. Variables analyzed were total acidity (Ac), Brix (Bx), total anthocyanins (TA), total polyphenols (TP). (A) Box-plots with median values and standard error bars. ANOVA results based on three biological replicates, with the different letters representing significant differences, Tukey HSD (*p*<0.05). (B) PCAs summarizing the phenotypic differences of the four selected clones for each analyzed year the greatest differences are driven by pH, TP and TA represented by PC1.

The RNA-seq analysis of berry pericarp of the four selected clonal accessions produced on average 45.7 million raw reads (6.7 Gb) per replicate (14x raw coverage), with 93.4% of the reads having base QC > 30 and an average GC content of 46% (Table S1C). After independent RNA-seq analysis, the sequenced reads aligned to 37,807 and 37,556 annotated transcripts in the Malbec-Pru and the Malbec-Mag haplophases, respectively. Despite this slight difference, the PCA based on global transcriptomic data showed that biological replicates of clone 595 were consistently differentiated from the other accessions (PC1 = 57%) (Figs 4A and B). According to their smaller phenotypic variation, the remaining three accessions appeared closer in the RNA-seq based PCA (PC2 = 17%) (Figs 4A and B). Differentiation of clone 595 was corroborated through clone pairwise comparisons, employing heatmaps of sample-to-sample distances (Fig. 4C and Fig. S6A to F). For all the described analyses the relations observed among the replicates of the four clonal accession was the same, using either of the haplophases as reference (Figs 4A-C and Fig S6A-F).

**Figure 4.**
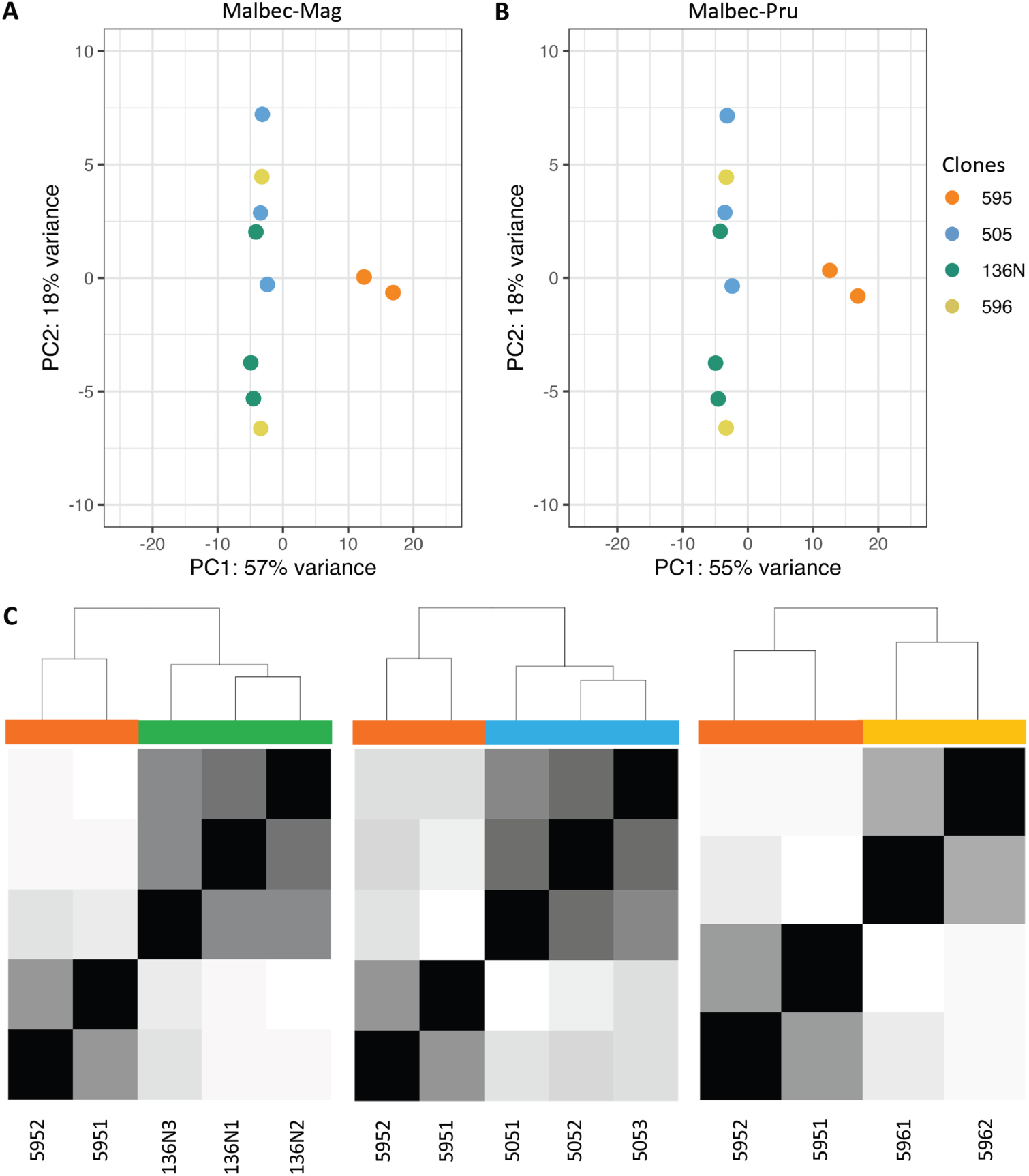
Principal components analysis based on global transcriptomic data based on total aligned transcripts to (A) Malbec-Mag and (B) Malbec-Pru. Both analyses showed consistent results, with the biological replicates of clone 595 differentiating from the rest based on PC1. The biological replicates of the other clones (136N, 505 and 506) differentiated less among each other based on PC2. (C) Pairwise comparison of sample-to-sample distance between 595 and the other three clones also showed clear differentiation. This analysis was performed with both haplophases and retrieved identical results; therefore, only Malbec-Mag results are shown.

To further investigate the differentially expressed genes (DEGs) that were driving differentiation of Malbec clone 595 from the others, pairwise comparisons were performed: 595vs136N, 595vs596 and 595vs505. For each comparison a very similar number of DEGs was detected with either haplophase as reference (Figs 5A and B; Table S6). The number of DEGs ranged between 392 and 722, with 595vs505 comparison producing the highest number of DEGs and 595vs596 the lowest. Hereafter, downregulated and upregulated DEGs are expressed in relative terms after comparing clone 595 to the other three. In all comparisons there were more downregulated than upregulated DEGs (Figs 5A and B; Table S6). In each comparison, on average 82% of the DEGs were detected for the orthologues of the two haplophases when using in parallel each haplophase as reference for the differential expression analysis (Fig. 5C). Moreover, 236 DEGs were consistently detected with the two haplophases in the three pairwise comparisons involving clone 595 (Fig. 5D). All these 236 DEGs showed the same direction of change using either of the two haplophases as reference (Table S6). On the other hand, from DEGs detected uniquely with one haplophase a higher number was detected with Malbec-Pru than with Malbec-Mag (Fig. 5C), whereas there were 11 DEGs consistently detected in the three comparisons only with Malbec-Pru and six DEGs detected uniquely with Malbec-Mag (Fig. 5D). On the other hand, for each pairwise comparison we analyzed the subcategory of DEGs that did not account for an annotated orthologous gene in the other haplophase (Table S6). We found that with Malbec-Mag an average of 7.4% of all the detected DEGs had no ortholog annotated in the Malbec-Pru haplophase (Table S6), this was significantly above (Fisher’s exact test *p*<0.05) of the total number of unmapped genes (5.4% = 1,915 genes) in the other haplophase. For Malbec-Pru the inverse scenario was observed, since an average of 3% of all the detected DEGs had no orthologue in Malbec-Mag (Table S6) and this was significantly below (Fisher’s exact test *p*<0.05) the total percentage of unmapped genes (2,296 genes = 6.4%). We also found that among DEGs with no ortholog annotated in the other haplophase, 20% in Malbec-Mag and 15% in Malbec-Pru were affected by structural variations (Tables S3C and S6), adding support on the putative hemizygous condition of these genes.

**Figure 5.**
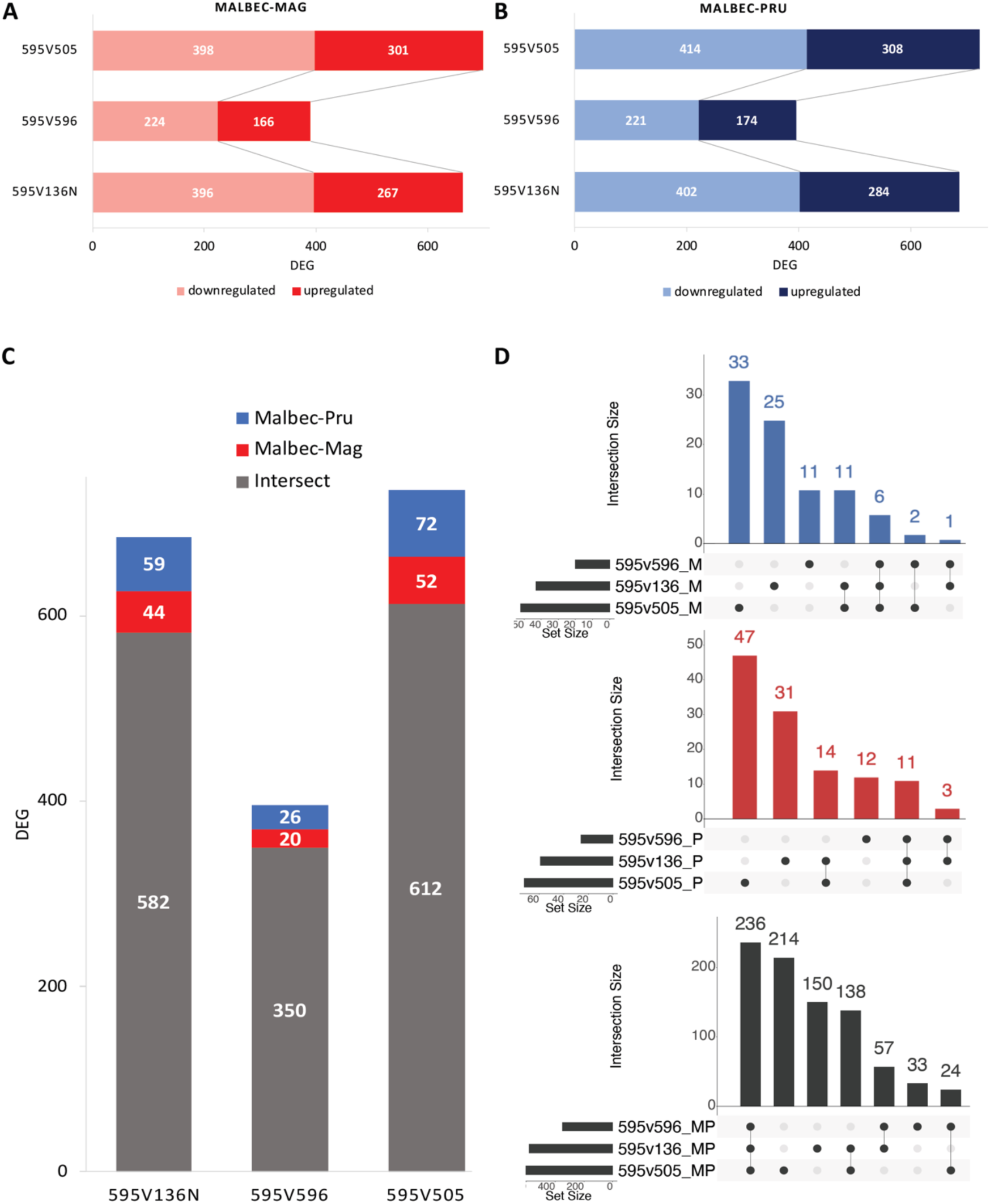
Total numbers of the statistically significant (*p*<0.01) differentially expressed genes (DEG). (A) and (B) Total number of DEGs detected in each clonal pairwise comparisons with Malbec-Mag and Malbec-Pru. Inside each bar downregulated and upregulated genes are distinguished based on the light and dark color code. (C) Intersection of the orthologous genes detected with both haplophases, for each clonal comparison. The grey portion of the bar represents the DEGs detected with both haplophases, the blue and red portions of the bar represent haplotype-specific DEGs detected with Malbec-Pru and Malbec-Mag, respectively. **d)** UpSet plots for the intersection among the three pairwise comparisons of the DEGs consistently detected with each haplophase, red bar-chart for Malbec-Mag, blue bar-chart for Malbec-Pru and black bar-chart for DEGs detected with both haplophases.

Gene Ontology (GO) enrichment analysis of biological processes (BPs) also produced similar results using either of the haplophases as reference, both for the enriched terms and the direction of change relative to clone 595. GO terms associated with ‘response to abiotic stress’, including ‘response to salt stress’ and ‘response to oxygen-containing compound’ where downregulated, whereas genes related to ‘secondary metabolism biosynthesis activity’ were upregulated in clone 595 (Figs 6A-C). Even though genes in the ‘response to abiotic stress’ GO term were mostly downregulated in 595, a number of DEGs in the same category were also upregulated, indicating for a possible deregulation of stress responses in berries of this clone (Fig. 6, Tables S6 and S8). In fact, a core of eight DEGs annotated in this category were consistently identified in all comparisons. Five out of the eight core DEGs in the ‘response to abiotic stress’ were consistently downregulated in 595 (Table S8), including: i) one gene (Mb_P_chr12g17550 | Mb_M_chr12g17200 | Vitvi12g02138) coding for a small heat shock protein (sHSP) assigned to the three enriched ‘response to abiotic stress’ child categories; ii) one gene coding for an ortholog of the DELLA protein GAI1 (Mb_P_chr01g05780 | Mb_M_chr01g05800 | Vitvi01g00446) assigned to ‘response to osmotic stress’ and ‘response to oxygen-containing compound’ child categories; and iii) three genes that coded for a SRM1 transcription factor (Mb_M_chr10g18440 | Mb_P_chr10g18110 | Vitvi10g01533), a Scarecrow-like protein 3 (Mb_P_chr06g12050 | Mb_M_chr06g12350 | Vitvi06g01133) and a C2H2-type domain-containing protein (Mb_M_chr01g09990 | Mb_P_chr01g10150 | Vitvi01g00845) assigned to the ‘response to oxygen-containing compound’ child category. On the other hand, three of the ‘response to abiotic stress’ core genes were consistently upregulated in 595 (Table S8). More precisely, two genes coding for sHSPs (Mb_M_chr13g05600 | Mb_P_chr13g05620 | Vitvi13g00491 and Mb_M_chr02g00360 | Mb_P_chr02g00330 | Vitvi02g00025), and one for a ZAT11 zinc finger protein (Mb_M_chr06g05190 | Mb_P_chr06g05230 | Vitvi06g01682). Overall, the function of the mentioned proteins is related to protein-folding homeostasis in response to stress ^38^ or regulation of transcription in the gibberellic acid signaling in response to osmotic stress ^39^.

**Figure 6.**
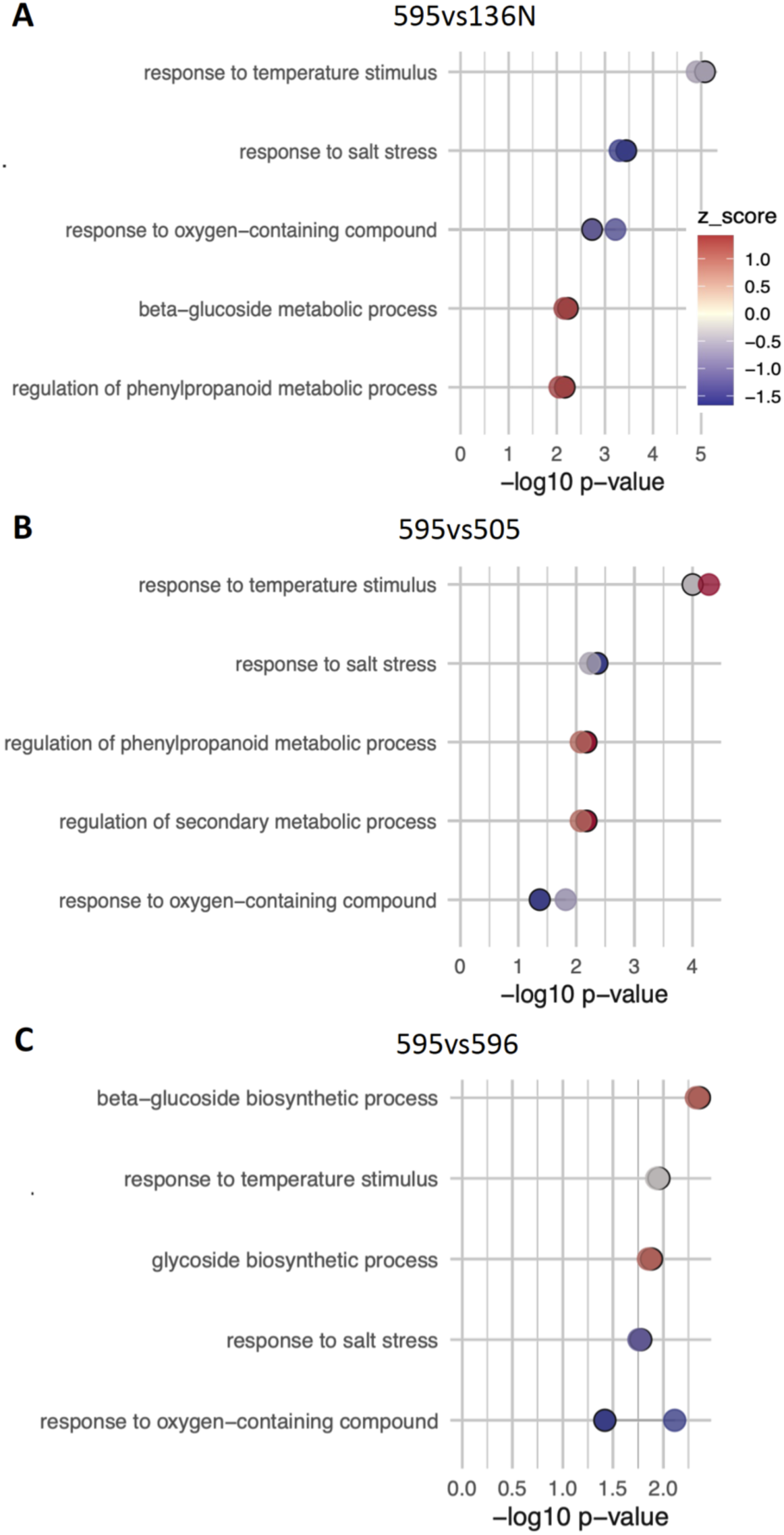
Biological processes enrichment analysis for the three pairwise comparisons (A) 595 vs 136N, (B) 595 vs 505 and (C) 595vs 596. For each process two dots are shown, indicating that the same process has been detected with both haplophases. Lighter dots represent the process detected with Malbec-Pru and darker dots with hard outline Malbec-Mag, in some cases dots are overlapped (and seen as a single dot) because the process has been detected with very similar *p-value*. Dots are colored based on its z-score value, blue indicates downregulation of the process, red is for upregulation and grey indicate values close to zero. The significance of the detected processes is expressed as the -Log10 p-value.

GO terms enriched with DEGs that were upregulated in clone 595 were associated with secondary metabolism, more precisely to the phenylpropanoid and beta-glucoside biosynthesis (Fig. 6A-C). A core of four DEGs assigned to these categories was consistently upregulated in the three pairwise comparisons and with both haplophases as reference (Tables S6 and S8), including i) two genes that were assigned to the ‘phenylpropanoid biosynthesis’ category, one coding for an Ultraviolet-B photoreceptor (UVR8-2) gene (Mb_M_chr13g04770 | Mb_P_chr13g04840 | Vitvi13g00350) and another coding an F-box/kelch repeat protein (Mb_M_chr05g19550 | Mb_P_chr05g19310 | Vitvi05g01703) (Table S8) and ii) two genes that were assigned to ‘beta-glucoside biosynthesis’ category, one annotated as putative UDP-glycosyltransferase in the reference gene catalogue (https://integrape.eu/resources/), (Mb_P_chr12g12840 | Mb_M_chr12g12580 | Vitvi12g01718) and a *VviUFGT* gene (Mb_P_chr16g02060 | Mb_M_chr16g01980 | Vitvi16g00156) encoding the UDP-glucose:anthocyanidin:flavonoid glucosyltransferase (UFGT) enzyme, that limits anthocyanin accumulation in grape berries ^40^. Another relevant gene in the anthocyanin biosynthetic pathway, *VvAOMT* (Mb_P_chr01g17010 | Mb_M_chr01g16440 | Vitvi01g04438), was upregulated in the 595vs136N and 595vs596, but not in the 595vs505 comparison (Table S6).

Among the hundreds of detected DEGs some have key regulatory functions that could be driving the described gene expression pattern. Furthermore, these genes appeared in the three comparisons with consistent change directions (Fig. 7). More precisely, two homeodomain-leucine zipper transcription factor (TF) encoding genes, *VviHB7* (Mb_P_chr15g12080 | Mb_M_chr15g11550 | Vitvi15g00912) and *VviHB12* (Mb_P_chr02g02730 | Mb_M_chr02g02810 | Vitvi02g00228), were upregulated in 595 in all comparisons (Fig. 7A). Orthologs of these genes play a role in stress-responsive co-expression networks mediated by abscisic acid (ABA) signaling ^41^. In fact, several ABA-related DEGs were consistently identified here in all comparisons, including the upregulation in Malbec 595 of the ABA biosynthesis gene *VviNCED3* (Mb_P_chr19g13140 | Mb_M_chr19g12630 | Vitvi19g01356), coding for a 9-*cis*-epoxycarotenoid dioxygenase that is limiting for ABA biosynthesis, and one PP2C gene (Mb_P_chr16g13850 | Mb_M_chr16g14410 | Vitvi16g01985); along with the downregulation of the *VviPYL7* ABA receptor (Mb_P_chr15g13250 | Mb_M_chr15g13250 | Vitvi15g00997) and an ABA-8 hydroxylase coding gene involved in ABA catabolism (Mb_P_chr18g15370 | Mb_M_chr18g15400 | Vitvi18g01304) (Fig. 7A). Another ABA receptor, *VviPYL1* (Mb_P_chr02g09580 | Mb_M_chr02g09810 | Vitvi02g00695), was upregulated in 595vs136 and 595vs505, whereas read counts were also higher in 595 when compared to 596 despite not being classified as DEG in the 595vs596 comparison (Fig. S7A and B). In addition, *VviMYB157* (Mb_P_chr17g10170 | Mb_M_chr17g09800 | Vitvi17g00822), a homolog of the *WEREWOLF* / *AtMYB66* TF, was upregulated in 595 in all comparisons (Fig. 7A). The allelic frequency of one SNV at *VviMYB157* suggested downregulation of the Magdeleine allele in the other three clones (Fig. S7C and D). Overall, the detected transcriptome variation during veraison correlated with the higher TA concentrations on mature berries of 595 (Figs 3, 6 and 7). Relations that are suggested but not proven are indicated with dashed lines. Genes with possible allele-specific overexpression in Malbec 595 suggesting for *cis*-acting regulatory somatic mutations are labelled with asterisks as candidate genes triggering all other downstream responses.

**Figure 7.**
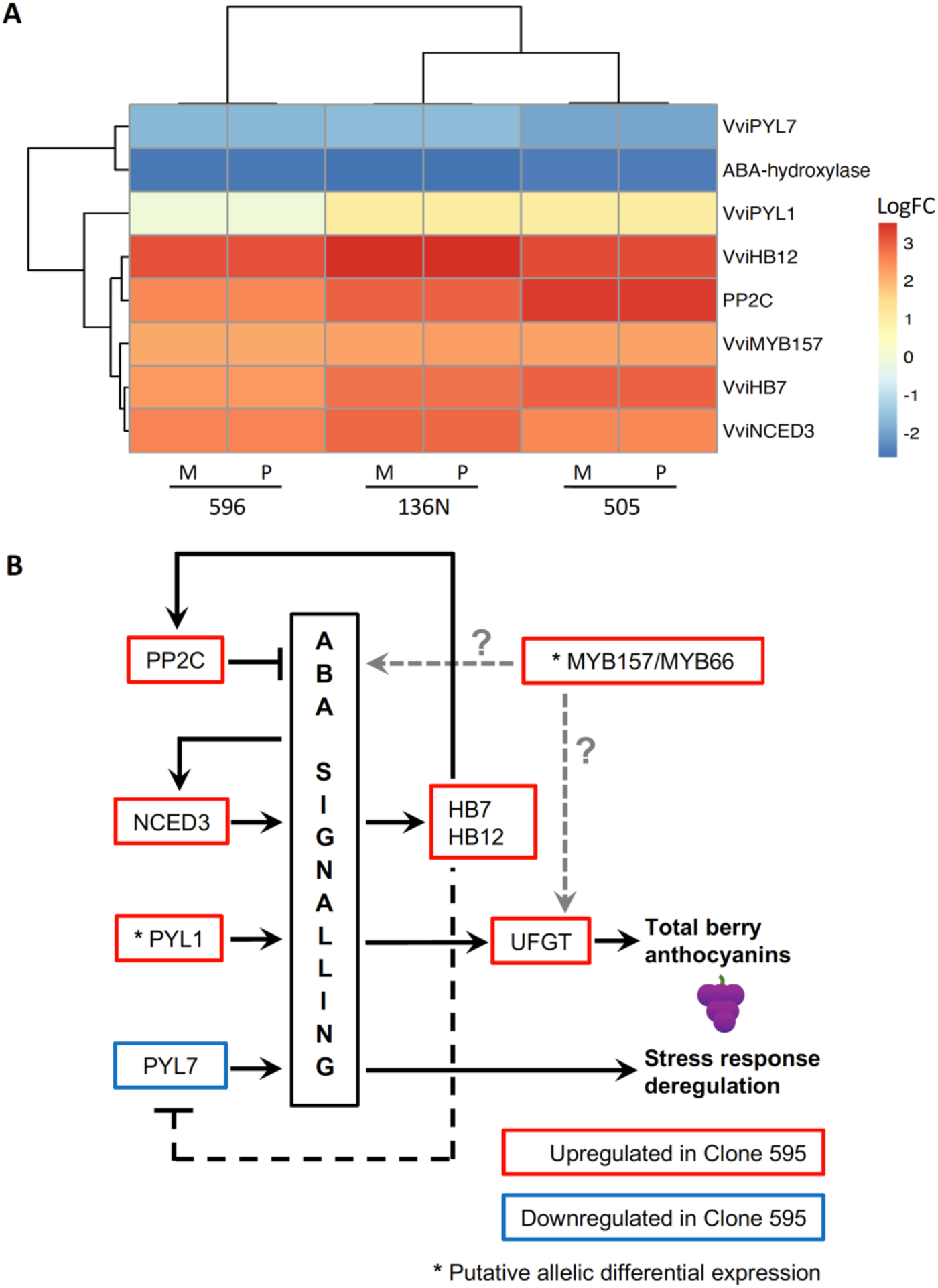
(A) Heatmap showing the change direction of DEGs with regulatory functions associated to the ABA signaling pathways. Log fold change of each gene is expressed in relative terms to 595. Results obtained with both haplophases (M = Malbec-Mag, P = Malbec-Pru) for the three pairwise comparisons are shown. (B) Proposed model based on the identified DEGs for the origin of the higher berry anthocyanin content in Malbec clone 595. Clonal variant phenotypes observed in 595 berries are indicated in bold. Genes upregulated in clone 595 compared to the other three are represented in red boxes, whereas the downregulated gene is in a blue box. Arrow and flat end lines indicate for positive and negative regulation.

## 4. Discussion

One of the main challenges for accurate genomic and transcriptomic clonal variation analyses in grapevine is to start with an appropriate reference genome for the cultivar of interest. Here we present the haplotype-resolved and annotated genome for cultivar Malbec, composed of the haploid complements inherited from its parental cultivars Prunelard and Magdeleine. We also employed Malbec diploid assembly as reference genome to perform a haplotype-aware evaluation of the transcriptomic differences underlying clonal phenotypic variation.

### The use of parental reads improved the phasing and evaluation of the assembly process

A haplotype-resolved genome assembly for Malbec was obtained in a trio-based approach applying prior knowledge of Malbec pedigree ^27^. Trio phasing and haplotype-aware deduplication yielded two haplophases that were close to the expected genome size of ∼475 Mbp and with low duplicated BUSCO genes (Table 1B). In this regard, our results were comparable to the obtained with the Merlot genome assembly, also based on trio binning ^19^. Trio binning approach differs from the applied for assemblies of other grapevine cultivars generated as pseudo-haplotypes, which exhibit inflated primary assemblies compared to the haplotigs ^6,9,11–14,42^. Nonetheless, recently developed bioinformatic tools help to improve the proper assignment of primary contigs to the two haplophases in those cases ^16^.

We analyzed the assembly consensus and the phasing accuracy based on *k-mers* from the parentals’ short-reads whole genome sequencing (Figs 1A and B). This had only been reported before in grapevine for the PN40024 reference genome ^2^. The estimated assembly accuracy for Malbec (QV > 40) indicates that error *k*-mers in the haplophase assemblies are < 0.00001%, which is slightly above the accuracy (QV=37.43) reported for the reference genome ^2^. Parental-specific *k-mers* (*hap*-mers) from short read sequencing of Malbec parents identified very few blocks that were apparently switched (<0.025%, Fig. 1 and Table 1C), in some cases these might be true homozygous regions in Malbec that were misassigned as switched. These results emphasize the importance of grapevine pedigree analysis and the value of using the parental reads, considering that some commercially relevant cultivars still lack this information despite the invested efforts (e.g. Nebbiolo) ^43^.

The diploid assembly also enabled a genome-wide study of the variation between the two Malbec haplotypes that showed ca. 25% of polymorphic regions (Fig. 2A and Table S3). The fraction of Malbec haplophases affected by SVs (12.1%) is much higher than in Zinfandel (6.5%) ^13^, but lower than in Nebbiolo (17.8%) ^9^ and close to Chardonnay (15.1%) ^6^, with the caveat that different SV evaluation methodologies were used, and that a thorough validation of reported SVs was performed only for Nebbiolo ^9^. In this direction, the percentage of genes overlapping with SVs between the two haplotypes in Malbec (29%, Table S3C) was in the range to the reported for Nebbiolo (26.2%). Overall, the fraction of the Malbec genome and genes affected by SVs is in agreement with the general observation that grapevine genomes can be unbalanced and that hemizygosity is widespread ^6^.

Almost half of Malbec genome consists of repetitive sequences and transposable elements (TE), with long terminal repeats (LTRs) representing the most abundant class of TEs (Fig. 1C and Table S4). This is in agreement with other grapevine cultivars ^9,18,42,44^, including the T2T assembly of the PN40024 genotype ^4^. Annotated gene number was similar in the two Malbec haplophases (Tables 2 and 3), and to the PN40024 v4 ^2^ and v5 (T2T) ^4^ versions of the reference genome. Nonetheless, close to 6% of genes did not have a counterpart mapping on the other haplophase. Furthermore, an enrichment of the resveratrol biosynthetic pathway in Malbec-Mag specific genes was observed (Fig. S3), suggesting for a putative higher contribution of this haplotype to a key secondary metabolite in grapevines, both from an industrial and biological perspective ^45^.

Orthologous genes between the two Malbec haplophases were mostly located in syntenic regions, whereas many genes assigned to the “Unknown” chromosome in PN40024 v4 could be placed on specific chromosomes across Malbec haplophases (Fig. 2B and Fig. S4). This suggests that the Malbec diploid assembly is improved in contiguity and representation of previously difficult-to-assemble regions. Moreover, when compared to the T2T HiFi sequencing-based assembly version of the PN40024 ^4^, Malbec haplophases have shown both strong structural correspondence and a congruent placement of most centromeres along the scaffolded pseudomolecules (Figs. 1C, S1C and S1D).

### Clone transcriptome variation in ripening berries could explain differences in berry anthocyanins content at harvest

We searched for consistent phenotypic and transcriptomic variation among Malbec clones that were cultivated in the same vineyard for over 20 years. Four clones were selected for analysis, based on their consistent variation in berry composition during 2018 and 2019 seasons (Fig. 3 and Fig S3A). According to a previous intra-cultivar genetic variation analysis, the four selected clones represent different derived genotypes from the same clonal genetic lineage (Group-Fr), with clones 595, 596, 136N and 505 belonging to genotypes I, B, H and E, respectively ^35^. Clone 595 had reproducible higher levels of total polyphenols and anthocyanins berry content at harvest (Fig. 3), whereas it also showed a clear transcriptome differentiation from the other clones (Figs 4 and 5). Functional enrichment of DEGs indicated for abiotic stress response deregulation in 595 compared to the other three clones (Fig. 6; Tables S6 and S8). On the other hand, phenylpropanoid and anthocyanin biosynthesis genes were consistently upregulated in 595 (Fig. 6; Tables S6 and S8), in agreement with the observed phenotypic differences. In our analysis, statically enriched processes (Fig. 6) correlated with the phenotype of the mature berry (Fig. 3). Similar relations were previously observed among Malbec clones diverging in berry anthocyanin content, when the transcriptional activity of post-veraison berries was studied ^32^.

Particularly, we identified ABA-related DEGs in Malbec clone 595 that might be major drivers of its global transcriptional profile and phenotype (Fig. 7A). ABA treatments have been reported to induce the gene *VviNCED3* and PP2C encoding genes and also to downregulate ABA receptor genes in different grapevine tissues including berry skin ^46^. Similarly, *VviNCED3* and an homolog of the ABA-inducible repressor PP2CA phosphatase ^47^ were upregulated in clone 595 compared to the other three, whereas the *VviPYL7* receptor gene was downregulated (Fig. 7A, Table S6). *VviHB7* and *VviHB12* class I HD-ZIP TF genes were also upregulated in 595. Their homologs in *Arabidopsis thaliana* have been implicated in a negative feedback loop induced by ABA and repressing ABA receptor genes ^48^. These expression patterns suggest that ABA over-accumulation or hypersensitivity could be a feature of clone 595. However, even though ABA usually represses *PYL* receptor genes ^46^, *VviPYL1* was upregulated in 595 (Figs 7A, S7 and Table S6). We hypothesize that somatic mutations in *VviPYL1* of clone 595 could trigger its over-expression and the hyperactivated ABA gene expression responses compared to other clones. To test this hypothesis, we inspected RNA-seq reads alignments for possible allele-specific over-expression of *VviPYL1* in 595, that could indicate the presence of a *cis*-acting somatic mutation in a regulatory region of this gene triggering its overexpression. However, both the RNA-seq reads and haplophase assembly comparison by SyRI support that Malbec is homozygous for *VviPYL1*, making it difficult to identify allele-specific differences among clones (Fig. S7). Therefore, further functional analyses are required to identify the underlying reasons explaining clone 595 differential response to ABA signaling.

Both the higher accumulation of anthocyanins and the differential expression of abiotic stress genes detected in clone 595 (Fig. 6, Tables S6 and S8) could be explained by an alteration in ABA homeostasis (Fig. 7B) ^49^. The overexpression of *VviUFGT* in 595 compared to the other three clones (Tables S6 and S8) could ultimately result in high berry anthocyanin content of this clone, since UFGT plays a critical role in the synthesis and accumulation of anthocyanins in grape berries ^50^. ABA treatment induces *VviUFGT* in grape berry skins, increasing its anthocyanin content ^46^. Therefore, ABA signal hyper-activation in 595 could potentially over-activate the flavonoid pathway leading to the higher phenolic character in 595 fruits (Fig. 7B). Transient overexpression of *VlPYL1* in grape berries induces ABA-responsive genes like NCEDs, but also *VviUFGT* ^51^, suggesting that the overexpression of *VviPYL1* in 595 could be an upstream trigger of both the ABA hypersensitivity and the high berry anthocyanin content in this clone (Fig. 7A-B). In addition, the putative allele-specific overexpression of *VvMYB157/MYB66* in 595 (Fig. S7) could be involved in the deregulation of the ABA signaling and the upregulation of the anthocyanin biosynthesis pathways characterizing this clonal line (Fig. 7B). *VvMYB157* might be particularly relevant in Malbec clonal anthocyanin content variation, since this cultivar is homozygous for *VvMAYBA1* and *VvMYBA2* loci and no evidence of differential gene expression was detected here (Fig. S7). A previous study of mature berries also reported the absence of variation for *VvMYBA1* expression among Malbec clones with different anthocyanins content ^32^. On the other hand, overexpression of a *MYB157/MYB66* homolog in the fruit skin of Russet apple somatic variants was associated with activation of ABA responses and the phenylpropanoid pathway ^52^. Our results warrant further functional genomic analyses to confirm the role of *VvMYB157* in the regulatory mechanism responsible of the high phenolic content found in Malbec clone 595.

### Phasing haplotypes helps to identify haplotype-specific gene expression

Analysis of transcriptomic data with either of the two Malbec haplophases as reference retrieved similar results, however some differences were observed. In this direction, 0.58% more transcripts aligned when Malbec-Pru was used as reference than with Malbec-Mag. This is in line with the fact that Malbec-Pru assembly turned 0.06% larger in size (Table 1B), accounting with 0.78% more protein coding annotated genes (Table 2) and 0.92% more transcripts predicted (Table 3), than Malbec-Mag. The analysis of cultivar Merlot haplotype-resolved genome also reported a similar difference, regarding annotated gene number between the two haplophases ^19^. In our analysis, the number of detected DEGs was 2.7% (±1.2% SD) higher with Malbec-Pru than with Malbec-Mag, on average for the three comparisons involving Malbec clone 595 (Fig. 5A and 5B). This observation is in agreement with the expected by chance based on the assembly and annotation features of both haplophases. Regarding the haplotype-specific DEGs, we found that six and 11 genes were consistently detected in all pairwise comparisons only with Malbec-Mag and Malbec-Pru (Fig. 5D), from which four and six had no ortholog in the other haplophase (Table S7), respectively. There is indeed a significant enrichment of Malbec-Mag putative hemizygous DEGs that were unmapped in the other haplophase, in contrast there was underrepresentation of Malbec-Pru DEGs that were unmapped in Malbec-Mag. Furthermore, two of the haplotype-specific DEGs detected only with Malbec-Pru coded for fruit development relevant functions. More precisely, an acyltransferase protein involved in fruit ripening (Mb_P_chr11g08440 | Vitvi11g04184) that was upregulated in clone 595, and an F-box like domain protein involved in the regulation of phenylpropanoids biosynthesis (Mb_P_chr08g18140 | Vitvi08g01787) that was downregulated (Table S7). Visual inspection of RNA-seq alignments on Malbec-Pru reference indicate putative hemizygosity for *Mb_P_chr11g08440*. In agreement, it has been shown that phased assemblies enhance the capture of molecular variation between clonal variants in highly heterozygous genomic regions, and that allele-specific expression is also associated to clonal phenotypic variation in other woody crops like citrus ^53^. On the other hand, two consistent heterozygous SNPs in *Mb_P_chr08g18140* indicate that Malbec might not be hemizygous for this locus, even though its ortholog was not predicted in Malbec-Mag haplophase (Fig. S7).

Overall, our findings are in agreement with the general observation that the genomes of grapevine cultivars are unbalanced and that hemizygosity is widespread ^6,10^. However, caution must be taken since miss-assembly and/or miss-annotation might be overestimating the reported putative hemizygosity. In this direction, the diploid assembly presented here could be useful not only to further corroborate putative allele-specific expression differences in highly heterozygous genomic regions^53^; but also for future introgression experiments, for which accounting with phased assemblies and transcript annotations has shown to be pivotal in grapevines ^54^.

## Conclusions

We have assembled the diploid genome of Malbec in two haplophases of the expected size and phase. Similar results were obtained in our RNA-seq comparison of Malbec clonal variants with either haplophase used as reference, nonetheless some haplotype-specific gene expression differences could be spotted thanks to the phased genome assembly. In this regard, genes with no ortholog annotated in the other haplophase of the Malbec assembly were enriched in relevant biological processes for grapevines, whereas a significant number of all the detected DEGs with Malbec-Mag had no ortholog in Malbec-Pru. SVs between the two assembled haplophases also suggested for hemizygosity of some DEGs contrasting between the studied clonal variants. Finally, we proposed that a somatic variant causing the hyperactivation of ABA signaling and the differential activity of a MYB transcription factor in a Malbec clone, might be responsible for its increased berry anthocyanin content.

## 5. Materials and Methods

### 6.1 Genome *de novo* assembly and annotation

#### Plant material

For genomic sequencing and assembly, samples were collected from the Malbec clone accession 136N, representing a clonal lineage with short propagation history in Argentina (i.e., Group-Fr) ^35^. Accession 136N is implanted at the Vivero Mercier Argentina experimental vineyard located at Lujan de Cuyo, Mendoza, Argentina (-33.09° S, -68.87° W). Samples of Malbec parental cultivars ^27^ were requested to the *Institut Français de la Vigne et du Vin* (VInopôle Sud-Ouest, France), corresponding to clone 1232 of Prunelard (reference number: 08-04-00004) and the unique available accession of Magdeleine Noire des Charentes (reference number: 08-04-00075).

#### DNA extractions, library preparation and genomic sequencing

Single node wood samples of the mentioned accessions were hydroponically grown, until the development of roots and shoots with young leaves. For PCR-free Illumina sequencing, DNA was obtained from roots for all three cultivars. For the parental cultivars solely roots were used, because this tissue derives from the L2 meristem cell layer (as gametes do in grapevine) and its genotype should be closer to the inherited by Malbec, in case chimeric somatic mutations existed. For Prunelard and Magdeleine, DNA extractions were performed with the DNeasy Plant mini kit (Qiagen). For Malbec, high molecular weight (HMW) DNA was obtained with the Nanobind plant nuclei kit (Circulomics-PacBio), either from roots or young leaves. Leaves were used for long read sequencing because it was the only tissue yielding the amount of HMW DNA needed for library preparation. In brief, nuclei were isolated from the source tissue according to Workman *et al*. ^55^. Isolated nuclei were the input for HMW DNA extraction with the Nanobind nuclei kit following manufacturer’s instructions. The HMW DNA was sheared with two strokes using a 26 G needle in a 1 mL syringe, with an average fragment size of 70 kb.

Library preparation and genomic sequencing were performed at the Max Planck Institute for Biology (Tübingen, Germany). A large insert PacBio gDNA library was prepared from the sheared HMW DNA of Malbec leaves using the SMRTbell Express Template Preparation Kit 2.0. The library was size-selected for >30 kb using BluePippin with a 0.75% agarose cassette (Sage Science) resulting in a 48 kb peak size final library. Five PacBio Sequel I SMRT cells were loaded with the same SMRTbell library and were sequenced with the PacBio Sequel I sequencer. Illumina PCR-free sequencing libraries were produced from gDNA of roots, using the NxSeq® AmpFREE Low DNA Library Kit (Lucigen) and sequenced to obtain paired-end short reads (150 bp), as described in Rabanal *et al.* ^56^.

#### Genome de novo assembly

The assembler Canu v1.8 ^57^ was employed for *de novo* genome assembly using the TrioBinning module ^15^. Here, parental-specific *k-mers* from the short-reads of the parental cultivars (Prunelard and Magdeleine) were used as template to split long reads of the child (Malbec) into the haploid complements. After adapters trimming with cutadapt v2.3 ^58^, Magdeleine and Prunelard short-reads were loaded into Canu assembler as *-haplotype1 and -haplotype2*, respectively, whereas Malbec PacBio CLR reads were loaded with option *-pacbio-raw*. The child haplotypes inherited by Malbec cultivar from each parent were separately assembled from the PacBio read partition, theoretically without the interference of inter-haplotype variation ^15^. Canu was run in all cases with option *genomeSize=490m*. No contaminant contig from non-plant genera or phyla was found from the draft assemblies by running blobtools v.1.1.1 ^59^, comparing to the non-redundant (nr) protein database using diamond v2.0.8.146 ^60^. The post-processing of the raw assembled contigs representing each haplotype of Malbec (hereafter: haplophases) involved a first round of polishing based on the long reads raw data using the ArrowGrid wrapper ^57,61^ of Arrow (http://github.com/PacificBiosciences/GenomicConsensus/) consensus framework, within the SMRT Analysis Software (http://github.com/PacificBiosciences/SMRT-Analysis). ArrowGrid was run on the primary assembly output of Canu with the corresponding haplotype-binned Malbec PacBio CLR reads for each haplophase, separately.

The assembly size metrics were tested along with the quality of the phasing process, to reduce haplotype switch errors ^15,62^. This was performed with Merqury v1.3 ^62^, based on the count of haplotype specific *k-mers* (i.e., *hap-mers*) from the short reads of Magdeleine and Prunelard as compared to Malbec short reads and the assembly *k-mers*. *Hap-mers* were used to determine phase blocks, defined as sets of markers inherited from the same parental haplotype ^62^. Each assembled haplophase genomic composition was tested using the default values, allowing 100 marker switches in a 20 kb bounded range (0.5% switch rate). Afterwards, complete contigs showing haplotype switch errors all along the contig and no-matches to the own haplotype in the Merqury output, were removed using a custom script (based on Unix/Linux *awk* and *grep* commands). Then, another haplotype-aware deduplication was performed based on minimap2 ^63^ alignments of one haplophase assembly to the other using *-x asm5* alignment option. Average similarity between haplotypes was computed for each contig from the *dv* value of the output PAF file. Candidate haplotype-switched contigs with high similarity to the other haplotype (average dv < 0.005) were extracted and used as reference to align the rest of contigs from the same haplophase with minimap2 (*-ax asm20 -secondary=no*). To check for possible duplicated contigs from that alignment, the coverage utility from samtools v1.10 ^64^ was run on the resulting *bam* file. High similarity contigs to the other haplotype were removed if they showed >70% bases covered, with depth ≥1 and mean depth of coverage >0.7 from the other contigs in the same haplophase. The resulting haplophases were further deduplicated with purge_dups v1.2.5 ^65^, read depth was used to remove duplicated contigs, by mapping the haplotype-binned Malbec PacBio reads to the corresponding haplophase assembly separately. The purge_dups command get_seqs was run with *-s* option to break contigs instead of adding Ns at internal breaks.

To perform miss-assembly corrections, RaGOO v1.1 ^66^ was used to split contigs that may generate spurious structural variations using the PN40024 12X.v2 ^3^ as reference, with options *-b -r 300000 -c 15000000 -d 30000000 -g 100 -s -C*. Afterwards, another round of assembly consensus correction based on long reads data was performed with ArrowGrid, in this case using all the Malbec PacBio CLR subreads to correct deduplicated contigs of the two haplophase assemblies concatenated. To overcome the high error rate of long reads, PCR-free Illumina short-reads of Malbec were used to polish the concatenated haplophases contigs, using Pilon v1.23 ^67^ with options *--fix snps,indels --minmq 3*. Then, haplophases were split according to their original contig IDs; and another round of haplotype-switch error detection was performed based on Merqury *hap-mer* analysis.

A final manual deduplication was conducted based on the gene content of the contigs spotted as haplotype switches by Merqury, and on BUSCO duplicated genes identified using the orthologue plant core genes database (eudicotyledons_odb10) and BUSCO v3.0.2 ^68^. When ≥2 BUSCO genes with different IDs were duplicated in the same order in another region of the same haplophase assembly, the duplicated copy spotted by Merqury as haplotype-switch error was considered as an artifactual haplotype duplication leaked to the wrong haplophase. Therefore, while the duplicated copy with correct haplotype assignment was kept in the final assembly, the duplicated and switched counterpart region was removed from the affected contig. This removal of duplicated and switched blocks was performed by running the *complement* and the *getfasta* applications of bedtools v2.27.1, using the contigs assembly (*fasta* file) and the duplicated coordinates (*bed* file). The boundaries of the coordinates of the duplicated regions to be removed were refined according to minimap2 alignments, with option *--secondary=no -x asm20 --MD* of each pair of duplicated contigs to the other haplophase contigs assembly. When no duplicated BUSCO genes were observed, then the contig spotted as switched by Merqury was consider as a likely homozygous region and was kept in the final assembly. In those cases, no corresponding haplotype switched block to be used for phase correction was present in the other haplophase assembly. After running the deduplication pipeline, another round of RaGOO miss-assembly correction was run as described before leading to only one break between chr2 and chr7 in Prunelard haplophase and none in Magdeleine haplophase.

RagTag v1.0.1 ^69^, using the PN40024 12X.v2 assembly as reference (including the 19 scaffolded chromosomes of grapevine, as well as the plastid and mitochondrial genomes), was employed for the final scaffolding of the deduplicated haplophases into the pseudo-molecules that represent the 19 chromosomes of grapevine haploid complement. The ragtag.py script was run with options *--remove-small -f 20000 -q 10 -d 500000 -i 0.5 -a 0.05 -s 0.3 -u --mm2-params "-x asm20*. Seqkit ^70^ was used to obtain assembly summary statistics of size and contiguity. SyRI v1.3 ^36^ was used to quantify the synteny and diversity between Malbec assembly haplophases. To run SyRI, the two haplophase assemblies were aligned with nucmer using options *-maxmatch -l 100 -c 500 -b 500* and delta-filter with options *-m -I 88 -l 100*. Nucmer and delta-filter were used from mummer v4.0.4 ^71^. Plots were produced with plotsr from SyRI. Completeness and duplication of contig and scaffolded assemblies of Malbec and of the PN40024 reference genome v4 and v5 versions was assessed running BUSCO v5.6.1^68^ on euk_genome_min mode, and using the eudicots_odb10 (Creation date: 2024-01-08) dataset as lineage and miniport ^72^ as gene predictor. Dot plots between the scaffolded Malbec haplophases and the 12Xv2 and v5 reference genome assembly versions were obtained running minidot on assembly-to-assembly alignments produced with minimap2 using *-x asm20* option.

#### Genome assembly annotation

Prior to gene annotation, EDTA v1.9.6 ^73^ was employed for the annotation of repeats and transposable elements (TEs), and to soft-mask the repeated regions. A repeat library was generated by running the EDTA.pl script with the options *--sensitive 1 --anno 1* and non-TE plant proteins were excluded from the repeat library. This analysis was run on the concatenated assembly (Malbec scaffolded haplophases) and separately, on the discarded contigs of each of the two haplophases that were not included in the final assembly after the deduplication pipeline. Firstly, blastx from blast v2.2.29 was run with options *-evalue 1e-10 -num_descriptions 10* to identify overlaps between the repeat library and a curated plant protein database obtained from the MAKER-P manual ^74^. Then, the overlaps were excluded from the EDTA repeat library by running ProtExcluder.pl ^74^. Finally, the curated repeat library was used to annotate TEs in the diploid assembly by running again EDTA.pl with options *--step anno --sensitive 1 --anno 1*.

For gene annotation a pipeline combining gene evidence obtained from different sources based on EVidenceModeler v1.1.1 ^75^ was implemented, using as input genome the soft-masked Malbec diploid assembly (concatenated and scaffolded haplophases). Here, *de novo* assembled transcripts, *ab initio* predictions and gene lift-over data were combined, and obtained as follows.

Transcriptome assembly was produced from an RNA-seq dataset obtained from berry pericarp of eight different Malbec clones and their biological replicates. Clones 225, 228, 59 and 53, were sampled during 2016 and employed solely for the annotation pipeline (Table S1C). For these samples total RNA extractions were performed from mature berries pericarp using a CTAB based protocol ^76^. Three to four berries were grounded with liquid nitrogen discarding the seeds, to obtain the desired amount of RNA (>3,5 ug). Afterwards, a column-based purification step was conducted with the kit *SV total RNA isolation System* (Promega). Extracted RNA integrity, concentration and purity were tested in 2% agarose gel and by spectrophotometry (AmpliQuant AQ-07). Samples were shipped to CRG facilities (Barcelona, Spain) for library preparation and sequencing. Library preparation was performed with a TruSeq Stranded mRNA LT Sample Prep Kit (Illumina) and included rRNA depletion using the Ribo-Zero kit (Illumina). Sequencing was performed with Illumina HiSeq 2500 and paired-end fragments of 125 bp were obtained. The four clones employed for transcripts *de novo* assembly, were also used for the clonal transcriptomic variability analysis (see that section for further details on total RNA obtention). Overall, a batch of 22 RNA-seq samples was employed for transcripts *de novo* assembly, adding ∼150 Gb of raw sequence (Table S1C). The Illumina sequencing adaptors and low quality reads were trimmed with TrimGalore v0.6.4 (Babraham Bioinformatics) that includes cutadapt v2.3 for read trimming and FastQC ^77^ for sequence quality checks. Two rounds of TrimGalore were run, the first to remove adaptors and low-quality reads with options *-q 20 --paired --length 30 --max_n 1 --trim-n* and the second to remove polyA tails and run the final QC with options *--paired --length 30 --polyA --fastqc*. For transcriptome assembly, the trimmed RNA-seq reads were aligned to the diploid Malbec genome assembly using HiSat2 v2.2.1 ^78^ with options *--rna-strandness RF --dta-cufflinks --max-intronlen 20000*. The obtained alignments (*bam* files) were merged into a single *bam* using the merge application of samtools v1.10 ^64^ and assembled into potential transcripts with StringTie v2.2.1 ^79^, using the options *-t -c 2 -f 0.1 -s 4.75 -M 0.95 -m 80 -j 2 -A --rf*. Starting from a genome-based transcript structure obtained with StringTie, the software TransDecoder v5.7.0 (Haas, BJ. https://github.com/TransDecoder/TransDecoder) was used to identify the likely coding sequences (CDs), adding them to the annotation files.

On the other hand, *ab initio* gene predictions were performed with Augustus v3.5.0 ^80^. Before running Augustus, Braker2 v2.1.6 ^81^ was run for the Malbec diploid assembly using as strand-aware hints the *bam* files produced by HiSat2 from the malbec RNA-seq data. This was performed to generate the retraining configuration file to be used as input in the *--species* option of Augustus. The EDTA soft-masked diploid assembly was used as input genome to run both Augustus and Braker2. To run Augustus, *extrinsic M.RM.E.T* information configuration was used as described in the *extrinsic.cfg* file (File S1) and the UTR prediction was inactivated *--UTR=off*. The hints input for Augustus was a *gff* file with exons annotated by TransDecoder from the StringTie transcriptome assembly of the merged Malbec RNA-seq (*gtf* file). Gene models resulting from Augustus were kept as the final *ab initio* gene predictions, but gene models were only kept for loci that showed some overlap with gene models that were also predicted in both, the final output of Braker2 and the output of Genemark that Braker2 uses for its retraining. The intersection between the three gene prediction version *gff* files was obtained by running a custom script based on *comm*, *grep* and the *intersect* -c command of bedtools v2.27.1 ^82^.

Also, a lift-over of the PN40024.v4 gene annotation to the Malbec haplophases assemblies was conducted with Liftoff v1.6.1 ^83^, with options*: -a 0.5 -s 0.5 -copies -sc 0.99 -exclude_partial -polish*. This included a final polishing step and transferred genes were only kept if similarity and coverage were >50% for the primary gene copy, whereas for secondary copies the similarity threshold was increased to 99%. The PN40024.v4 gene annotations of the improved grapevine reference genome assembly ^2^ were retrieved on 15.11.2021, from the Integrape Cost Action CA17111 portal (https://integrape.eu).

The weighed consensus gene models for Malbec diploid and unmasked genome assembly were obtained with EVidenceModeler, using the evidence obtained from the described sources. The final combination of evidence inputs and weights was selected after testing more than 50 EVidenceModeler runs and comparing the obtained gene models to the deeply curated PN40024.v4 ^2^. For EVidenceModeler, the RNA-seq transcriptome assembly obtained with StringTie was used as TRANSCRIPT input with the maximum weight = 4, the lifted-over of PN40024.v4 reference annotations obtained with Liftoff was loaded as OTHER PREDICTION (weight = 3), and the gene model produced by Augustus was used as AB INITIO PREDICTION with the minimum weight = 1.

Finally, the *annotation update* function of PASA v2.4.1 ^75,84^ was used to polish exon boundaries and to update the 5’ and 3’ UTRs to the genes structures and alternative isoform annotations, previously predicted by EVidenceModeler. Here, the input transcripts were the ones assembled with StringTie from the Malbec RNA-seq samples and the transcripts annotated in the primary (REF) and alternative (ALT) haplotypes of the PN40024.v4. All these transcripts were cleaned from polyA, low quality or vector sequences using the *seqclean* script of PASA. Cleaned transcripts were aligned to the Malbec diploid assembly with Blat ^85^ and Gmap ^86^ aligners using the *Launch_PASA_pipeline.pl* script with *– transcribed_is_aligned_orient* option and the following options in the alignAssembly.config file: validate_alignments_in_db.dbi:--MIN_PERCENT_ALIGNED=90; validate_alignments_in_db.dbi:--MIN_AVG_PER_ID=95; subcluster_builder.dbi:-m=50; run_spliced_aligners.pl:-N=5; validate_alignments_in_db.dbi:--MAX_INTRON_LENGTH=25000. Two consecutive rounds of PASA updates from the aligned clean transcripts were run using again the *Launch_PASA_pipeline.pl* script, by loading firstly the EVidenceModeler annotations and secondly the updated annotations resulting from the first round. Genes were renamed according to assembly chromosome coordinates using the script retrieved from https://github.com/andreaminio/AnnotationPipeline-EVM_based-DClab ^87^.

To perform QC at each annotation step, the general metrics of the predicted gene models were computed using the *agat_sp_statistics.pl* script from AGAT v0.8.0 ^88^. Transcripts and translated proteins were extracted from *gff* annotation files with Gffread v0.11.7 ^89^ and the general stats were computed with Seqkit v0.12.0 ^70^. Completeness and duplication levels for the annotated protein sets were obtained running BUSCO v5.6.1 ^68^(on protein mode), using the eudicots_odb10 (Creation date: 2024-01-08) dataset lineage.

Functional annotation to predict gene ontology (GO) classes and gene functional description (DE) was performed with Panzzer2 ^90^, using as input the *fasta* files containing all the protein sequences of each haplophase. The correspondence between the obtained gene model predictions for each haplophase, PN40024.v4 and VCostv3 annotations was performed using the proteins amino acid sequences, with OrthoFinder ^91^. Afterwards, GENESPACE ^92^ was used to evaluate the order and location of the predicted gene models along the scaffolded pseudomolecules of the assembled haplophases, compared to the PN40024.v4 as the most complete gene predictions for the reference genome (Table 3B). The co-location of annotated genes with genome variation between the two assembled haplophases of Malbec was performed using the *intersect* command of bedtools v2.27.1 to compare the final gene predictions of each haplophase to the genome variation output of the comparison of the two haplophases produced by SyRI.

Genes overlapping with each variation feature assigned by SyRI were analyzed for enrichment in Biological Process GO terms using g:Profiler ^93^, with the same settings described below for the functional enrichment analysis of DEGs.

To annotate centromeres and telomeres, a search for the 107 nt monomer (CEN107= AGTACCGAAAAAGGGTCGAATCAGTGTGAGTACCGAAAAATGGTAGAATCCGGGCGAGTACCGGGAAAAGGTAGAATCCGTGCGAGTATCGAAAAACTGTCCGGGCG) of centromeric repeats and the 7 nt monomer (TTTAGGG) of telomeric repeats was performed ^94^. EDTA v1.9.6 was used to annotate centromeric repeats on the Malbec diploid assembly by running the EDTA.pl command with *--sensitive* 1 *--anno* 1 parameters from a repeat library containing only the CEN107 monomer. Telomeric repeat positions were identified on the diploid assembly using BLAST v2.2.29+ ^95^. Blast alignment was run with oligonucleotide query parameters (*blastn* -task *blastn-short*) using as query two consecutive copies of the 7-mer monomer. Finally, the density of all the annotated elements was plotted along the assembled pseudomolecules using ggplot2, by taking the coordinates of the start position of each annotated element as well as of variants detected between Malbec haplophases.

### 6.2. Analysis of the phenotypic and transcriptomic clonal variation

#### Clonal phenotypic variation analysis

A survey focused on berry composition variation was performed on 27 Malbec clones, with three biological replicates, resulting in 81 plants. All analyzed plants were located in the same plot at Vivero Mercier Argentina experimental vineyard (Luján de Cuyo, Mendoza, Argentina), implanted since 2002. Therefore, the analyzed plants have been exposed to the same cultural treatments and environmental conditions ever since. Phenotypic measurements were conducted during 2018 on mature berries. For each biological replicate a random sample of 50 berries was obtained and the pulp was separated from the skin. The pulp was used for analytical measures (pH, total acidity and total soluble solids) and the skin was used for spectrometry analyses (total anthocyanins and total polyphenols). The pH was directly measured with a Denver Instrument (UB-10, pH/mV Meter). Total acidity (expressed in g/L of tartaric acid) was measured through acid-base titration with 0.1 N NaOH (pH = 8.2), using 5 mL of juice diluted with 10 mL of distilled H2O and containing bromothymol blue (0.04%). Total soluble solids were measured in wt/wt of sucrose, expressed in Brix degrees (Bx), using an ATAGO refractometer (PAL-1). The skins were dried and weighed to perform total polyphenols extraction using a ethanol and water solution (12:88, v/v), containing 5 g/L of tartaric acid, according to Muñoz *et al.*^32^. Total polyphenols (TP) and total anthocyanins (TA) were measured with a UV-VIS spectrophotometer (Spectrum SP-2000UV) at wavelengths of 280 and 520 nm respectively, and expressed in AU/g of skin. The same procedures were repeated for the phenotypic analyses in 2019, only for the clonal accessions selected for transcriptomic analyses.

#### Plant material selection for clonal diversity transcriptomic experiments

Four Malbec clones (595, 596, 136N and 505) were selected after a ranking-based procedure, based on phenotypic data obtained for the 27 clones in 2018. The selected samples combined high, mid and low values for the surveyed traits, except for sugar content trait where the selected clones had either high or low values, aiming to represent as much as possible of the observed variation (Fig. S4B). ANOVAs and Tukey HSD tests were performed for pairwise comparisons between the selected clones, to test for significant differences among the analyzed traits (*p*<0.05). Berries used for total RNA extraction were sampled during *veraison* stage (switch point from berry development to ripening, when 75% of berries were colored); a transcriptionally active stage for genes involved in secondary metabolism pathways ^96^. Sampling was performed previous to 2019 harvest, within the same day (01.15.2019) and during morning time (between 8-10 a.m.); the collected berries were kept in liquid nitrogen until final storage in ultra-freezer (-80°C).

#### Total RNA extraction, library preparation and sequencing

Total RNA extractions and quality checks were performed as previously described (annotation section). RNA samples were shipped to Novogene (Beijing, China) for sequencing. At arrival, samples were tested before library preparation and all passed the QC performed for quantitation (NanoDrop, ThermoFisher), degradation (agarose gel) and integrity (Agilent 2100, Bioanalyzer systems). Library preparation was performed with a TruSeq Stranded mRNA LT Sample Prep Kit (Illumina) and included rRNA depletion using the Ribo-Zero kit (Illumina). All samples passed the library QC performed for concentration (Qubit 2.0, ThermoFisher), insert size (Agilent 2100) and library effective concentration (qPCR). An Illumina HiSeq 2500 instrument was used for sequencing, to obtain 150 bp paired-end reads.

#### Differentially expressed genes and functional enrichment analyses

The base quality of the raw reads was checked with FastQC ^77^. Trimming of sequencing adaptors and low quality sequences was performed with Trimmomatic ^97^. rRNA and tRNA sequences were removed from the trimmed Illumina reads using Bowtie2 ^98^ aligner, by mapping the reads to a specific grapevine database ^99^. All forthcoming bioinformatic procedures were performed following a haplotype-aware scheme, using separately and in parallel both Malbec haplophases as reference genomes. Alignment of the trimmed reads was conducted with STAR ^100^, the assembly of the aligned reads with StringTie ^79^, and GffCompare ^89^ was used for extraction and annotation of transcripts. The read counting of the aligned *bam* files was performed with FeatureCount ^101^, according to Chialva *et al*. ^102^.

Heatmaps and PCAs displaying the global gene expression were built in R, based on the regularized log transformation of the normalized counts (rld), obtained with DESeq2 ^103^. To detect differentially expressed genes (DEG) with DESeq2 ^103^, pairwise comparisons among the four clones were performed. The threshold to consider a gene to be differentially expressed (DEG) was set to a *p-*value<0.01 (FDR-adjusted) and an absolute value of log2 fold-change |LFC|≥1. Functional enrichment analyses were conducted using the list of DEGs with g:Profiler2 ^93^, to find statistically significant gene ontology (GO) terms related to Biological Processes (BP) and Molecular Functions (MF). For all GO enrichment analysis the g:GOSt function of g:Profiler was implemented using the PN40024.v4 functional annotation as reference. Aiming to compare DEGs and BPs identified with each of Malbec haplophases, the orthologue genes obtained with Orthofinder were employed. For all performed GO terms enrichment analyses, only manually curated annotations of experimental and computational studies were considered, and electronically inferred annotations were dismissed to avoid sporous results. This was performed based on a *p-*value <0.05 (g:SCS-corrected for multiple comparisons ^93^). The aggregate scores and z-scores where obtained and graphically represented with the R package GeneTonic ^104^. Z-scores summarize the direction of change based on the LFC values of the gene set involved in the enriched process. Positive z-scores (color red) indicate that a BP GO term is upregulated, whereas negative z-scores (color blue) indicate downregulation of a BP ^104^. GeneTonic function *gs_overlap* = 0.9 was employed to summarize redundant GO terms and *gs_summarize_overview_pair* was employed to make dumbbell plots, to visualize simultaneously the enriched GO terms detected with both haplophases. The web tools UpSetR (https://gehlenborglab.shinyapps.io/upsetr/) and jVenn ^105^ were employed to perform upset plots and Venn diagrams for DEGs pairwise comparisons.

## 6. Acknowledgements

This work was supported by Agencia Nacional de Promoción Científica y Tecnológica (ANPCyT): PICT2018-02381, MINCyT/ANPCyT (FONTAR)-CDTI IBEROGEN; CONICET (Bilateral PCB-II, CONICET-CSIC; MINECO BIO2017-86375-R, and the project PID2020-120183RB-I00 funded by MCIN/AEI/10.13039/501100011033), and the Max Planck Society. L.C. was supported by a CONICET travel grant for a short-term staying at the Max Planck (Tübingen). We are grateful to Ilja Bezrukov for answering bioinformatics question, and to Silvina van Houten for collaboration with field work. We thank Olivier Yobregat (Institut Français de la Vigne et du Vin, IFV) for providing plant material from Malbec parental cultivars. This project received funding from the European Union’s Horizon 2020 research and innovation programme under the Marie Sklodowska-Curie grant agreement No 797460. This study benefited from the networking activities and resources produced within the COST action Integrape (CA17111) and COST innovators grant Grapedia (IG17111).

## 7. Author contributions

L.C. coordinated the project and wrote the article. Produced and analyzed phenotypic, transcriptomic and genomic data. P.C.B. laboratory work for genomic sequencing and designed the bioinformatic pipelines for genome assembly and annotation, analyzed genomic data, and global data interpretation. C.M. phenotypic and transcriptomic diversity analysis. L.B. field and laboratory work for phenotypic diversity analysis. D.B. and C.S. project design, provided insight and access to the analyzed clonal accessions. W.T. bioinformatic analysis, set-up and maintenance of genome browse at IBAM-CONICET web-page. C.L. laboratory work for genomic data obtention. S.G.T., J.I. and J.M.M.Z. project design for genomic and transcriptomic experiments. C.R., J.I. and J.M.M.Z. obtained parental cultivars plant material and provided resources for whole DNA extractions. D.W. provided resources, facilities and advice for genomic laboratory work and bioinformatic analyses. D.L. designed and coordinated the entire project, performed transcriptomic analysis. All authors read and improved this manuscript

## 8. Data availability statement

Malbec-Mag and Malbec-Pru assemblies are available at the NCBI public repository under BioProjects: PRJNA1036636 and PRJNA1036637, respectively. All raw data is available at NCBI under BioProject: PRJNA1037531, including genomic Illumina short-reads obtained for cultivars Malbec, Prunelard and Magdeleine, PacBio long reads for Malbec and a batch of 22 RNA-Seq from eight Malbec clones and replicates. Malbec annotated assemblies are also available for visualization and gene browse at two public repositories: IBAM-CONICET (http://ibam.mendoza-conicet.gob.ar/resources/genome-visualizer/) and Gramene (https://www.gramene.org/). A gene correspondence between Malbec-Pru and Malbec-Mag annotated genes, alongside with PN40024.v4 and VCost.v3 annotations is provided, therefore gene queries can be performed using any of the mentioned formats.

## 10. Supplementary data

Supplementary data is available at Horticulture Research online.

