## Supplementary material for "Diploid genome assembly of the Malbec grapevine cultivar enables haplotype-aware analysis of transcriptomic differences underlying clonal phenotypic variation": File S1

# extrinsic information configuration file for AUGUSTUS
# 
# include with --extrinsicCfgFile=filename
# date: 15.4.2015
# Mario Stanke


# source of extrinsic information:
# M manual anchor (required)
# P protein database hit
# E EST/cDNA database hit
# C combined est/protein database hit
# D Dialign
# R retroposed genes
# T transMapped refSeqs
# W wiggle track coverage info from RNA-Seq

[SOURCES]
M RM E T

#
# individual_liability: Only unsatisfiable hints are disregarded. By default this flag is not set
# and the whole hint group is disregarded when one hint in it is unsatisfiable.
# 1group1gene: Try to predict a single gene that covers all hints of a given group. This is relevant for
# hint groups with gaps, e.g. when two ESTs, say 5' and 3', from the same clone align nearby.
#
[SOURCE-PARAMETERS]


#   feature        bonus         malus   gradelevelcolumns
#		r+/r-
#
# the gradelevel colums have the following format for each source
# sourcecharacter numscoreclasses boundary    ...  boundary    gradequot  ...  gradequot
# 
# The exonpart malus of .995 means a weak penalty factor for every predicted exonic base that is not supported by any exonpart hints. 

[GENERAL]
      start      1          1  M    1  1e+100  RM  1     1    E 1    1    T 1  1
       stop      1          1  M    1  1e+100  RM  1     1    E 1    1    T 1  1
       exon      1          1  M    1  1e+100  RM  1     1    E 1  1e4    T 1  5e3
     intron      1          1  M    1  1e+100  RM  1     1    E 1  1e4    T 1  5e3
        CDS      1          1  M    1  1e+100  RM  1     1    E 1    1    T 1  1
        UTR      1          1  M    1  1e+100  RM  1     1    E 1    1    T 1  1
nonexonpart      1          1  M    1  1e+100  RM  1     1.15 E 1    1    T 1  1
#
# Explanation: see original extrinsic.cfg file
#
