## Supplementary material for "Diploid genome assembly of the Malbec grapevine cultivar enables haplotype-aware analysis of transcriptomic differences underlying clonal phenotypic variation": Fig. S1

**Supplementary figure 1**. Minidot plots showing the strong structural correspondence between the 19 scaffolded pseudomolecules of Malbec haplophases and the grapevines reference genome. (A) and (B) shows the comparison between Malbec haplophases with reference version used here for the scaffolding process (PN40024_12Xv2). (C) and (D) shows the comparisons between Malbec haplophases with the latest version available of the grapevine reference assembly PN40024.v5 (T2T).

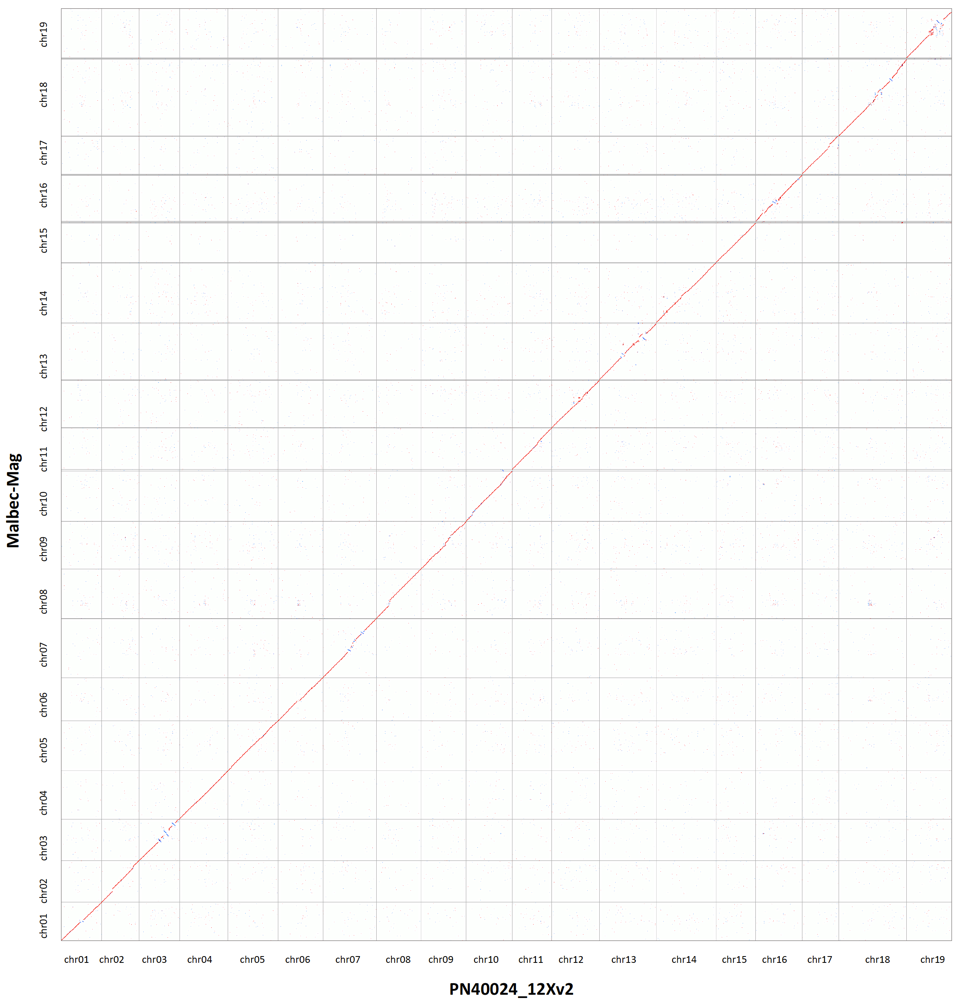
A)

B)

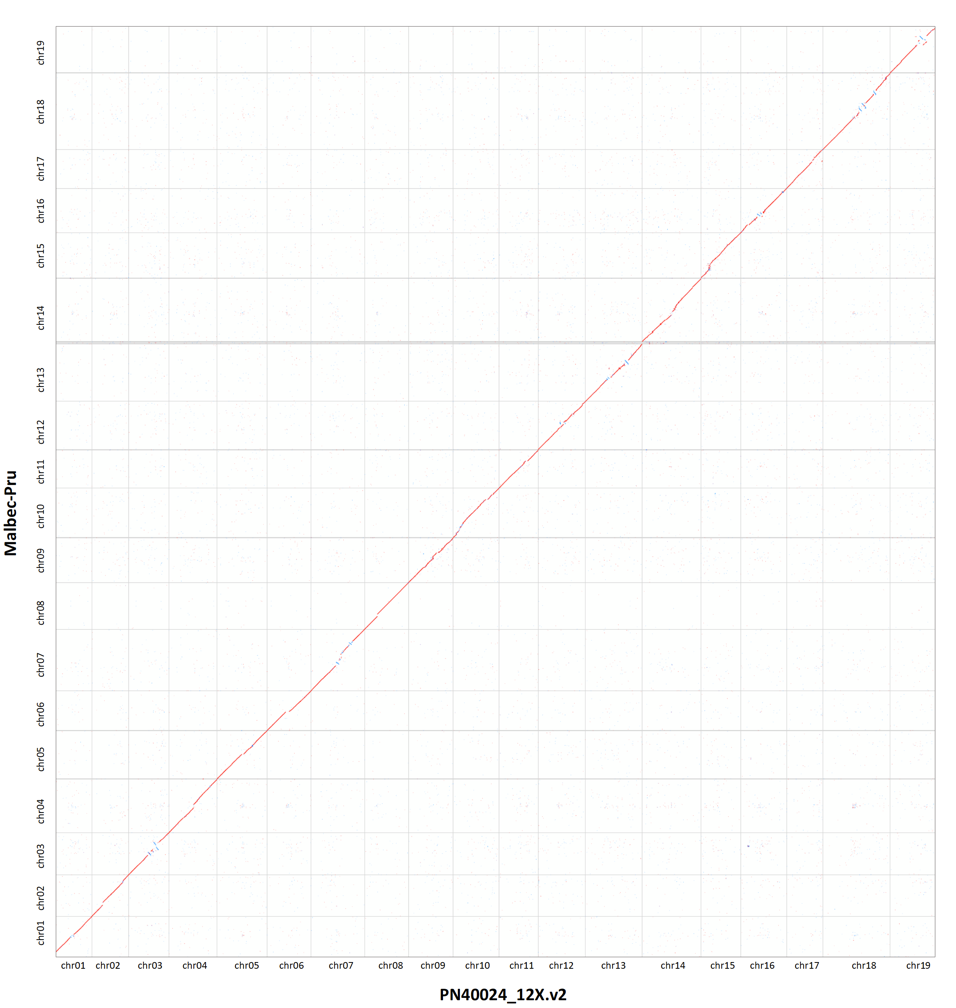

C)

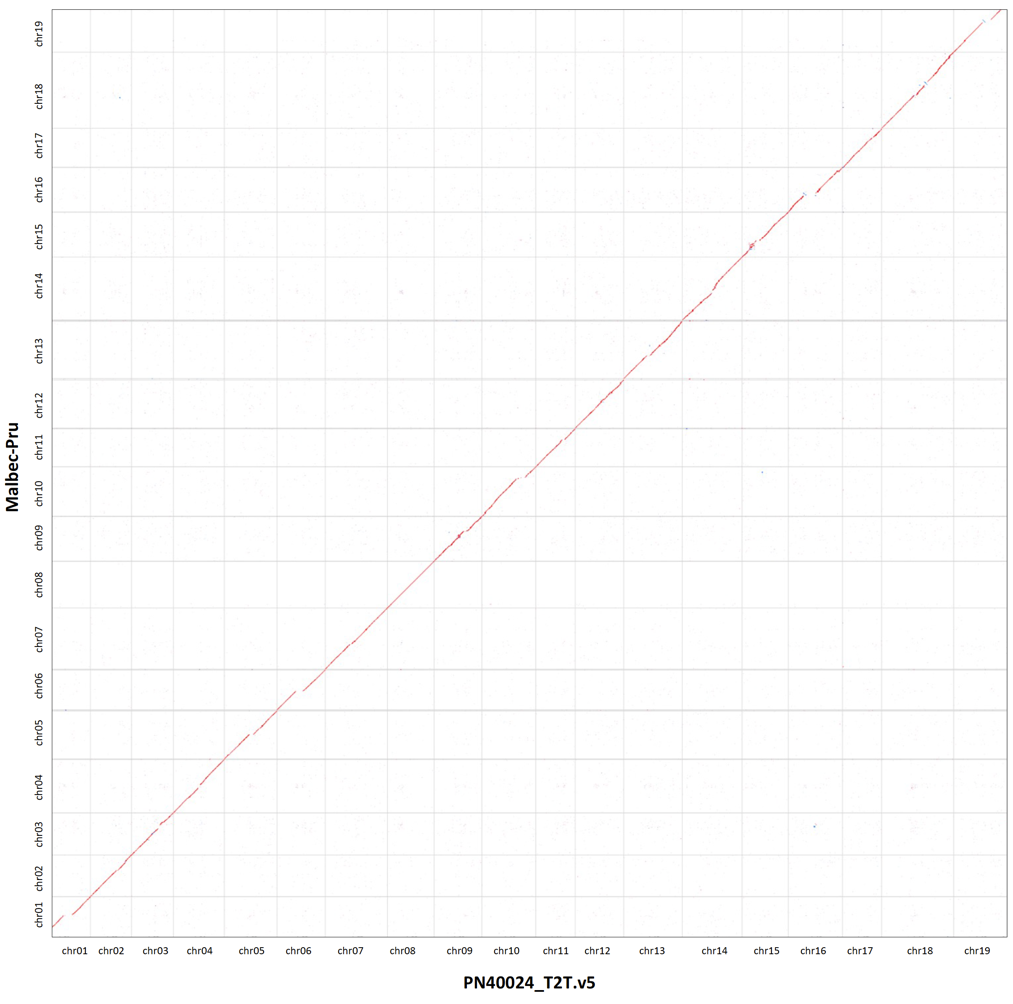

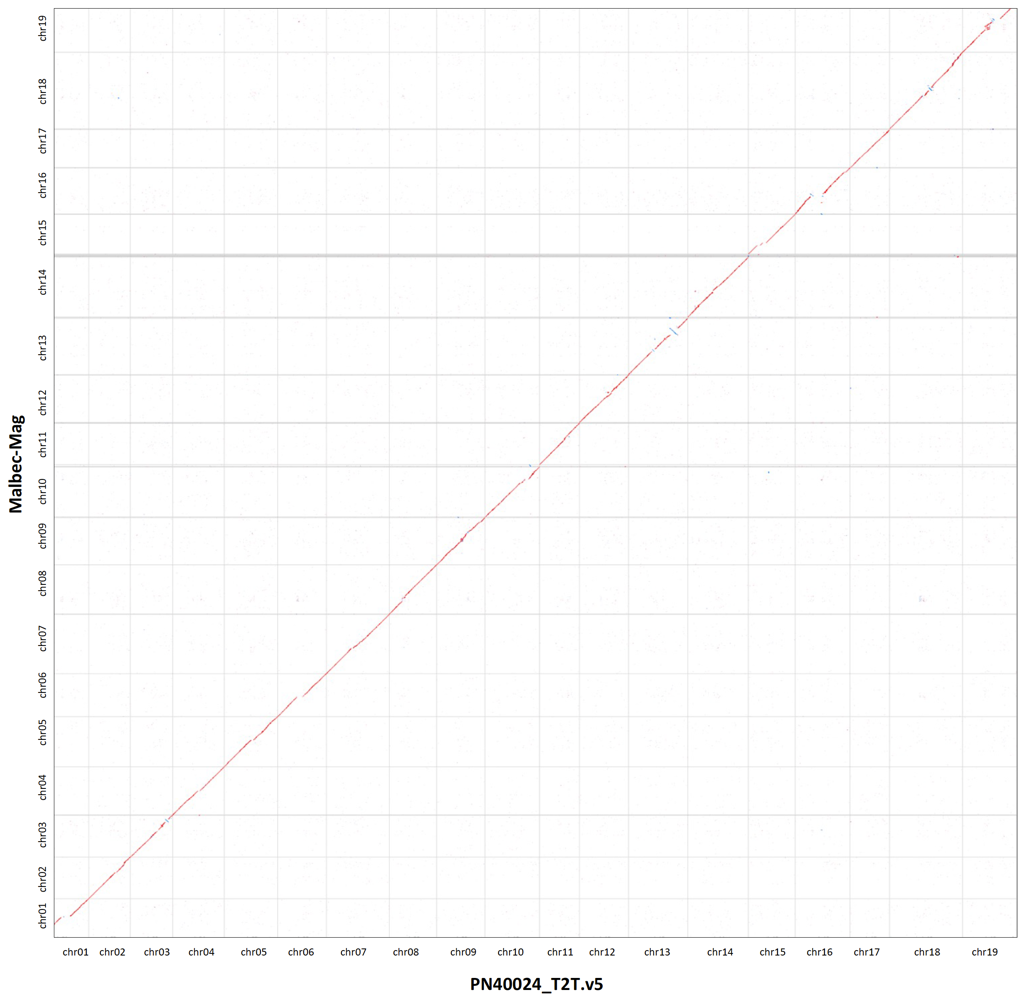
D)
