## Supplementary material for "Diploid genome assembly of the Malbec grapevine cultivar enables haplotype-aware analysis of transcriptomic differences underlying clonal phenotypic variation": Fig. S2

**Supplementary Figure 2**. Contiguity (N50) of the obtained contigs and scaffolds, compared to the phased blocks for **(A)** Malbec-Mag and **(B)** Malbec-Pru haplophases. Phased blocks contiguity was estimated after mapping the parental reads k-mers (hap-mers) to the respective assembly.

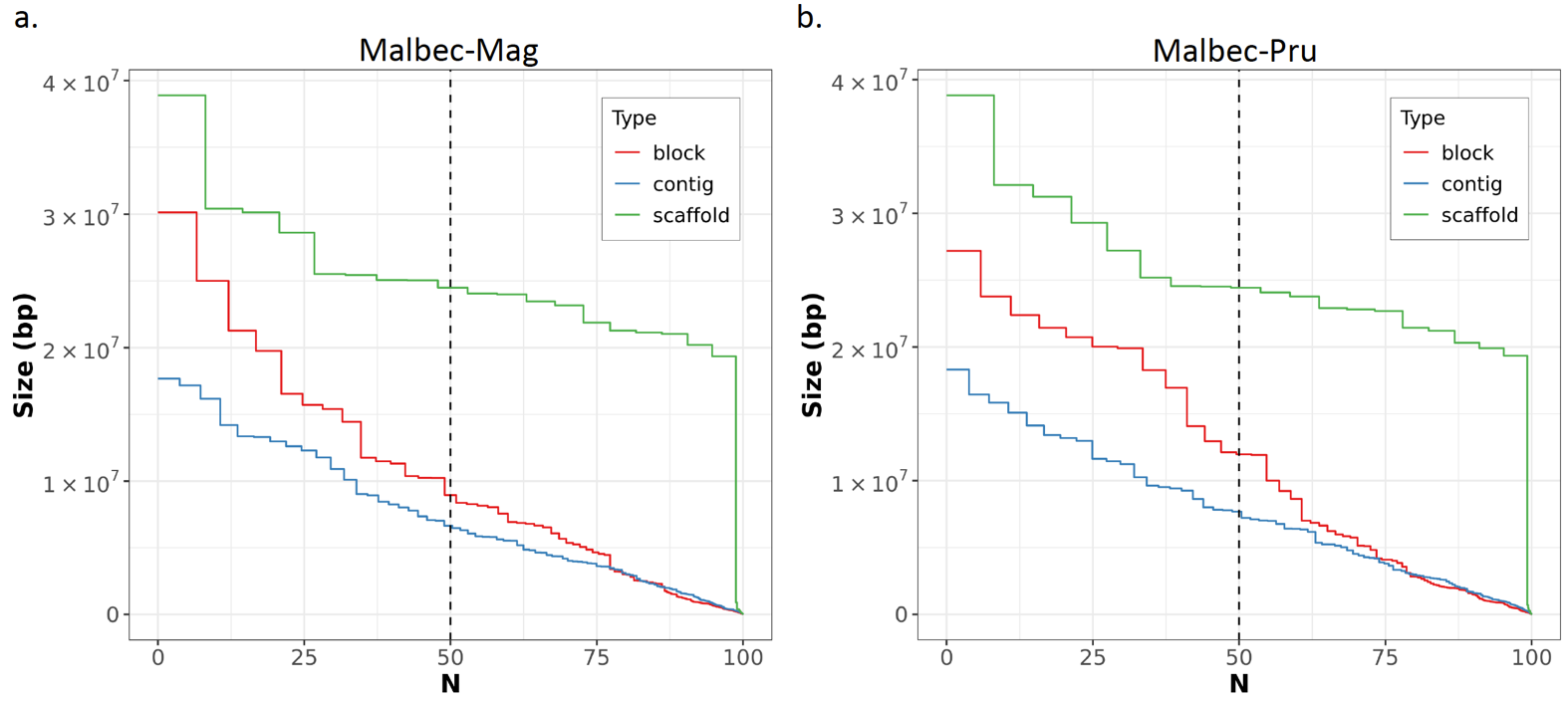
