## Supplementary material for "Diploid genome assembly of the Malbec grapevine cultivar enables haplotype-aware analysis of transcriptomic differences underlying clonal phenotypic variation": Fig. S3

**Supplementary Figure 3**. GO enrichment analysis performed with g:Profiler of the unmapped orthologous genes identified uniquely in **(a)** Malbec-Mag and **(b)** Malbec-Pru haplophases. Manhattan-like plots are shown with the main driver GO terms highlighted in each plot, the size of the circles is relative to the number of genes representing the term. Also, tables with details of the enriched term are provided below plots.

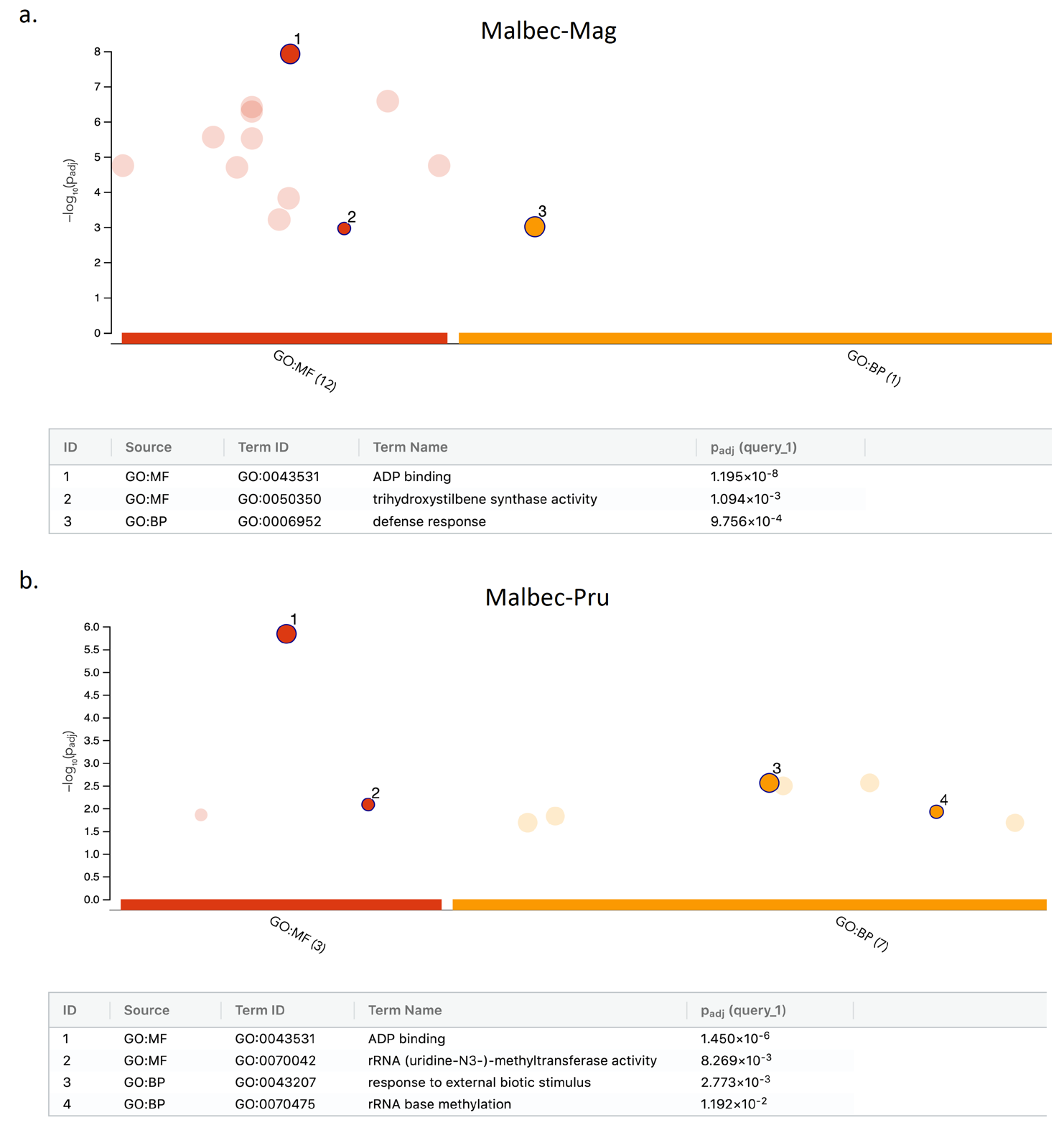
