## Supplementary material for "Diploid genome assembly of the Malbec grapevine cultivar enables haplotype-aware analysis of transcriptomic differences underlying clonal phenotypic variation": Fig. S4

**Supplementary Figure 4**. Dotplots representing the gene orthogroups synteny between, (A) Malbec-Mag and (B) Malbec-Pru with the grapevine reference genome (PN40024.v4). Most orthogroups are aligned to the diagonal indicating strong synteny. The red rectangles highlight orthogroups located in the ‘Unknown’ chromosome of the reference genome, that exhibited orthogroups counterparts assigned to different chromosomes across Malbec assemblies.

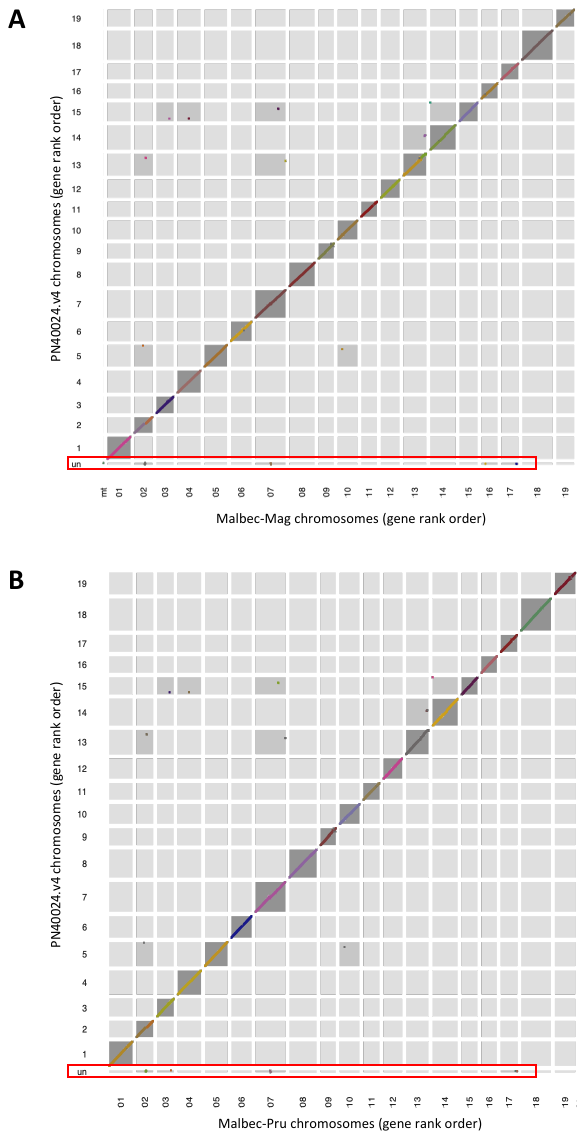
