## Supplementary material for "Diploid genome assembly of the Malbec grapevine cultivar enables haplotype-aware analysis of transcriptomic differences underlying clonal phenotypic variation": Fig. S5

**Supplementary Figure 5**. (A) Principal components analysis including the 27 analyzed Malbec clones evaluated in 2018 harvest season. Berries composition was measured at harvest, total anthocyanins (TA) total polyphenols (TP) and pH were influencing differentiation on PC1 (46.2%). Differences along PC2 (27.26%) were mainly driven by Total acidity (Ac) and sugar content (Bx). (B) Table summarizing the phenotypic traits of the selected four accessions to perform RNA-seq analysis. (C) Timeline summarizing the sampling points, source material and objective of the sampling.

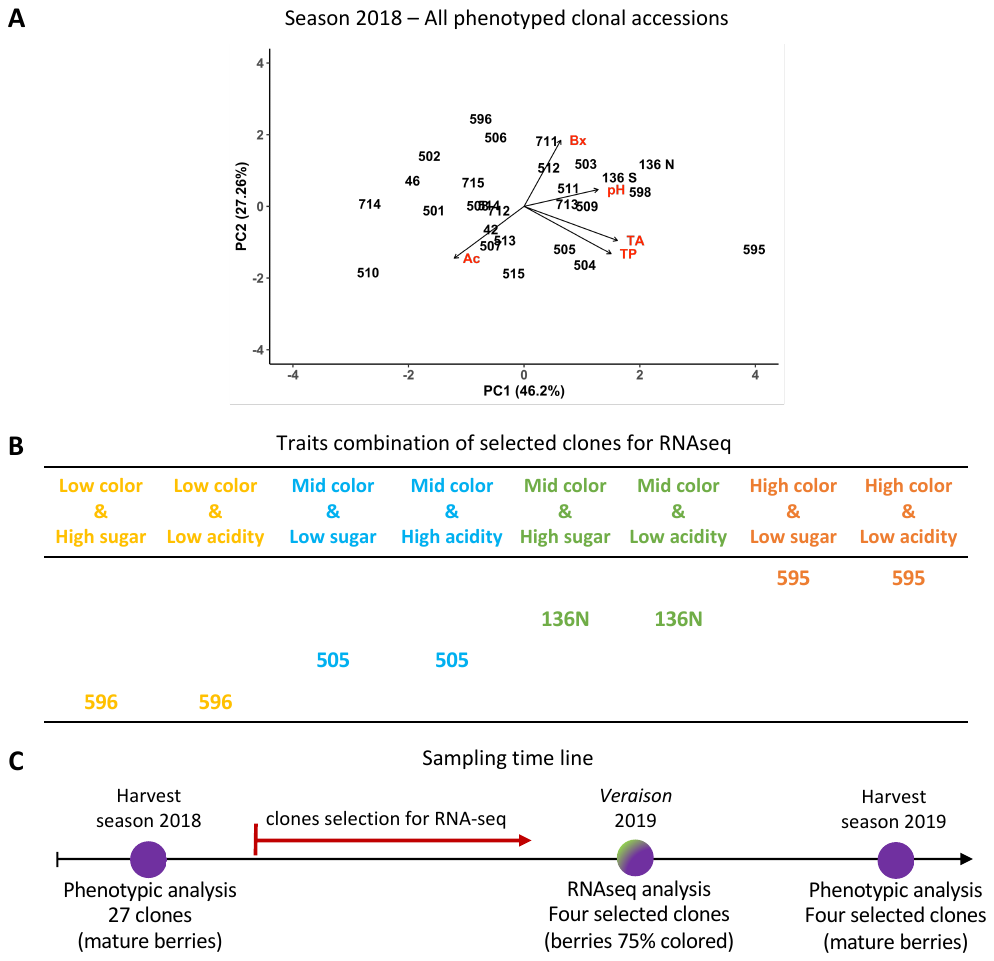
