## Supplementary material for "Diploid genome assembly of the Malbec grapevine cultivar enables haplotype-aware analysis of transcriptomic differences underlying clonal phenotypic variation": Fig. S6

**Supplementary Figure 6.** Pairwise comparisons based on global transcriptomic sample-to-sample distance. Pairwise comparisons were performed with Malbec-Mag (A to C) and Malbec-Pru (D to F). The obtained results were identical with either haplophase, in all cases Malbec clone accession 595 differentiated from the other accessions.

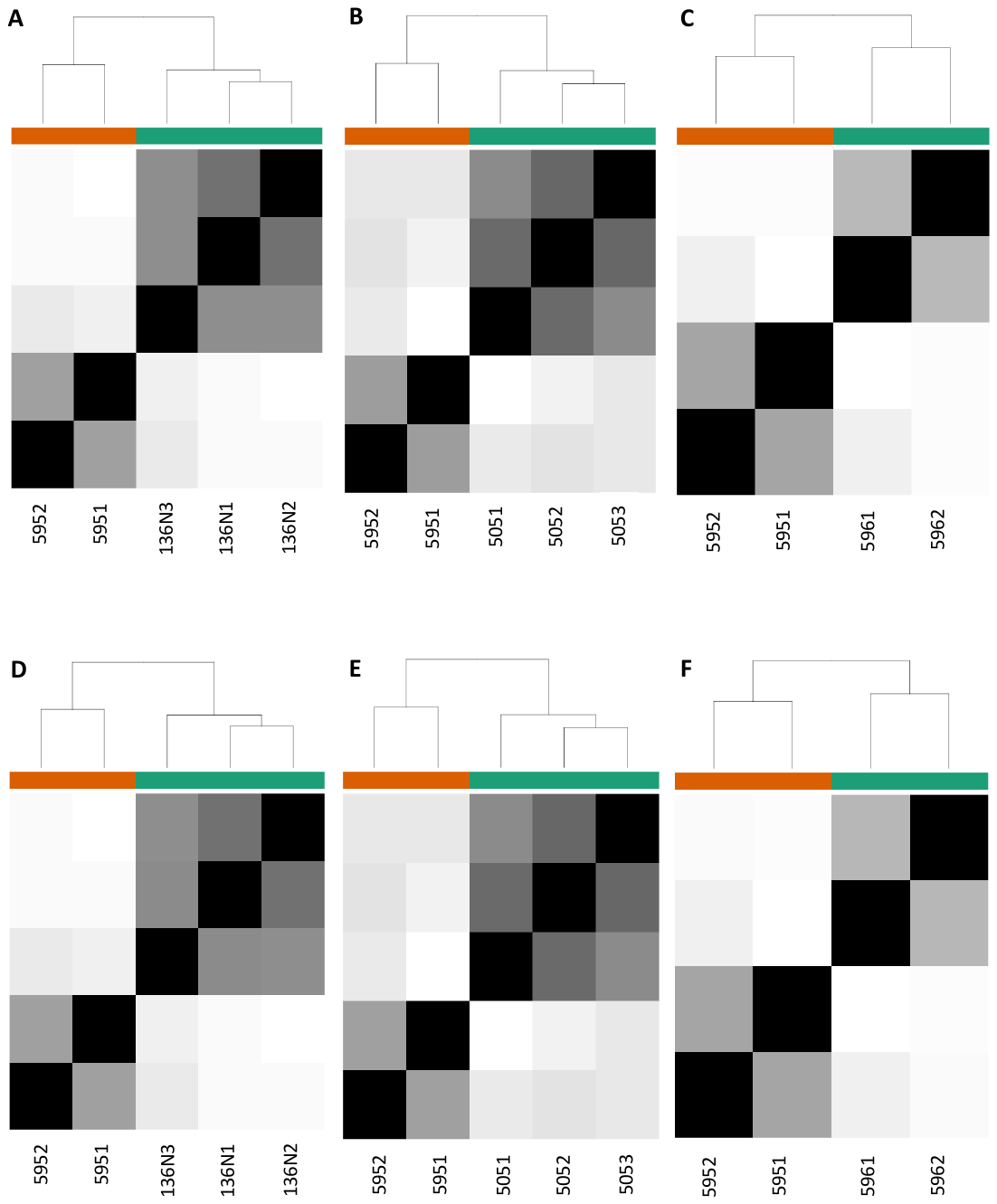
