## Supplementary material for "Diploid genome assembly of the Malbec grapevine cultivar enables haplotype-aware analysis of transcriptomic differences underlying clonal phenotypic variation": Fig. S7

**Supplementary Figure 7**. IGV screenshots to visualize the transcriptomic data alignment for the DEGs of interest. Each screenshot shows the biological replicates for the four analyzed accessions aligned to the assembly, also in the bottom of the screenshot the gff file showing the gene structure and bed file indicating for possible SNPs across the analyzed positions. (A) and (B) show bam files of PYL1 aligned to Malbec-Mag and Malbec-Pru respectively to show that PYL1 has more count reads than the other clones, even when it was not called as DEG versus 596. (C) and (D) show bam files for the MYB157 gene. The reference and alternative frequencies allele are shown, to visualize the putative allele specific expression of the Magdeleine-inherited in 136N, 505 and 596. (E) and (F) bam files for MYAB1 and MYBA2 aligned to the both haplophases and for all the analyzed accessions, to show their homozygous status in Malbec without differential expression. (G) bam files for the two putative hemizygous loci for Malbec-Pru, showing SNPs that discard that condition.

A

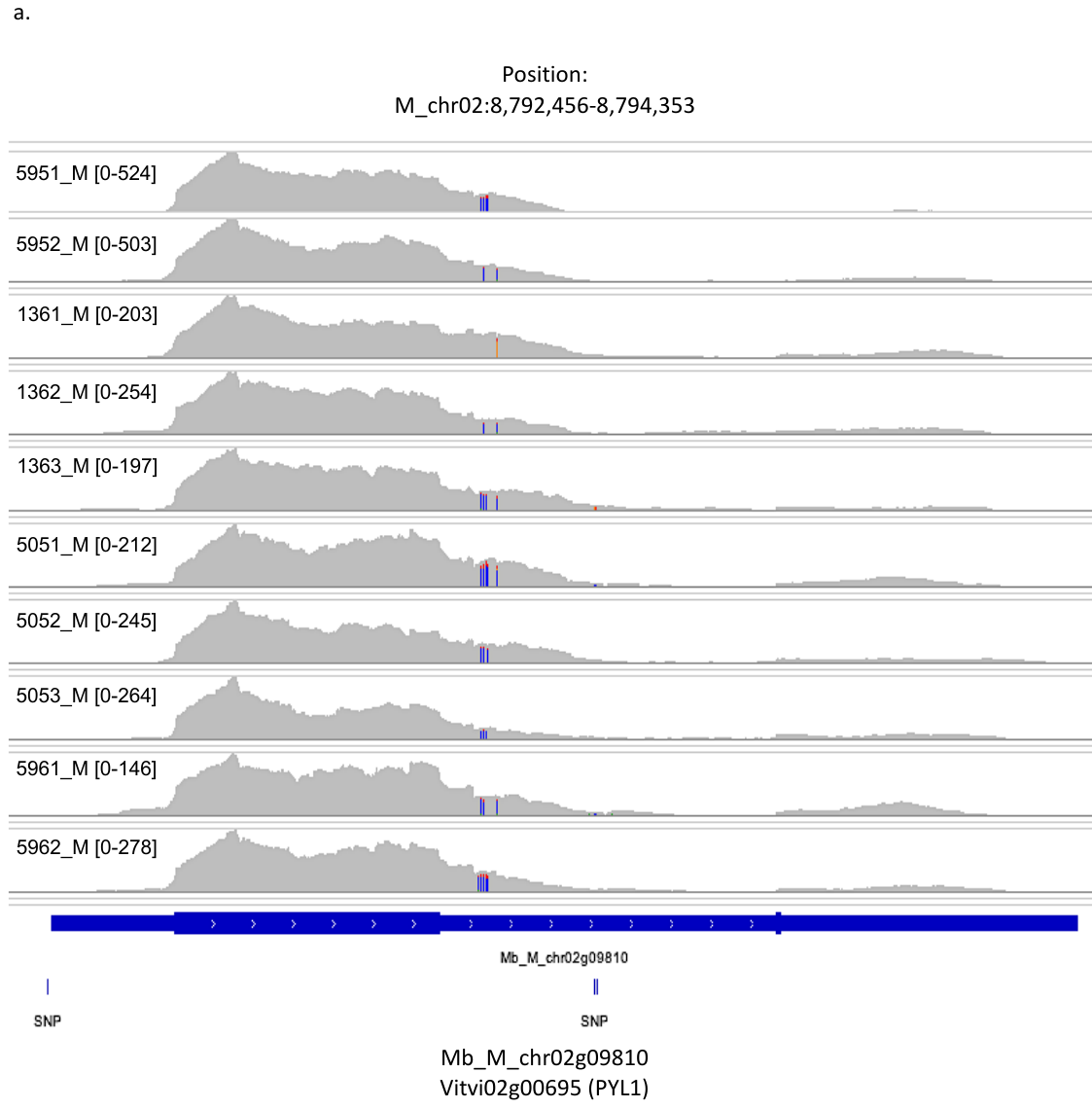

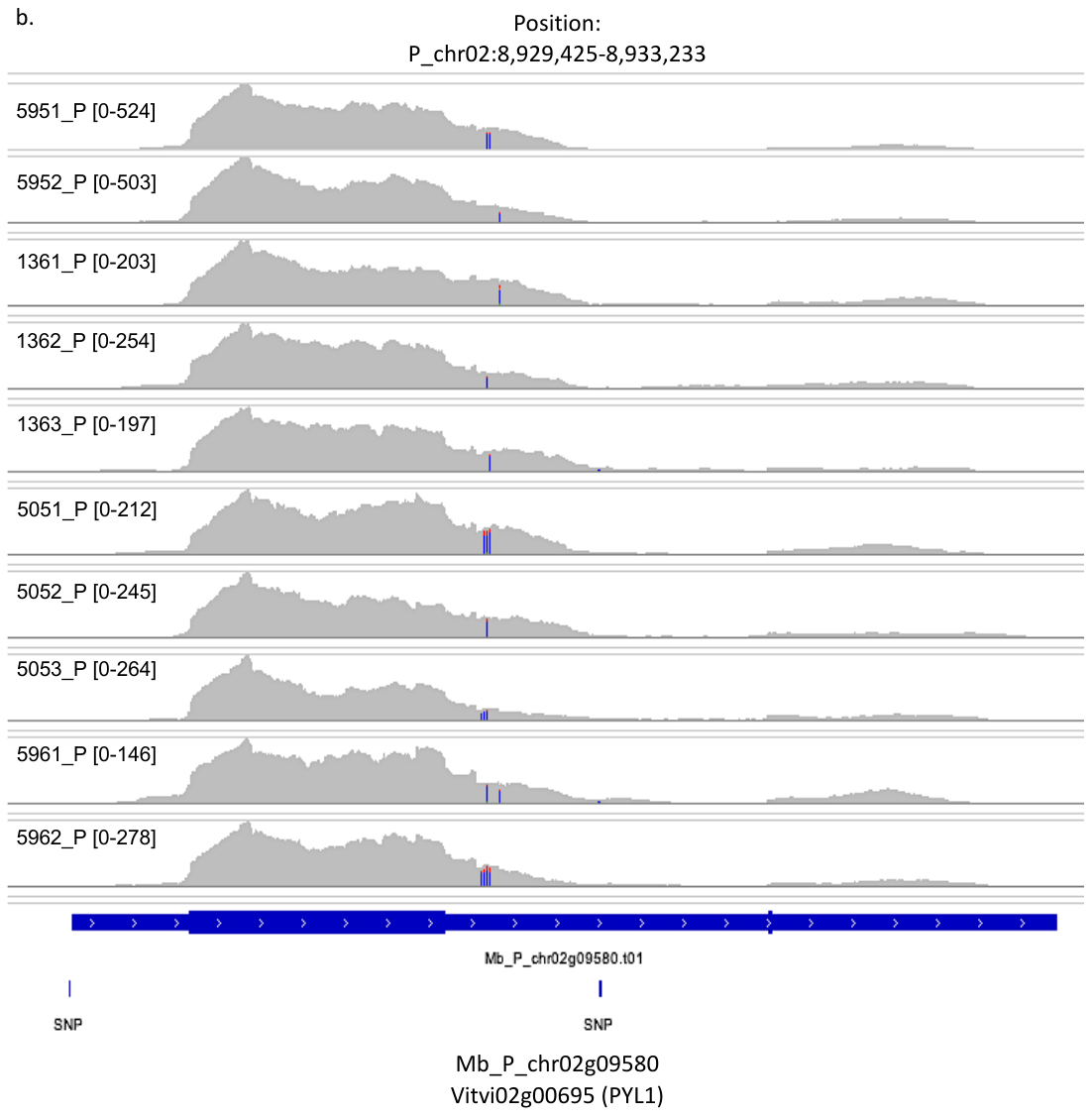

C.

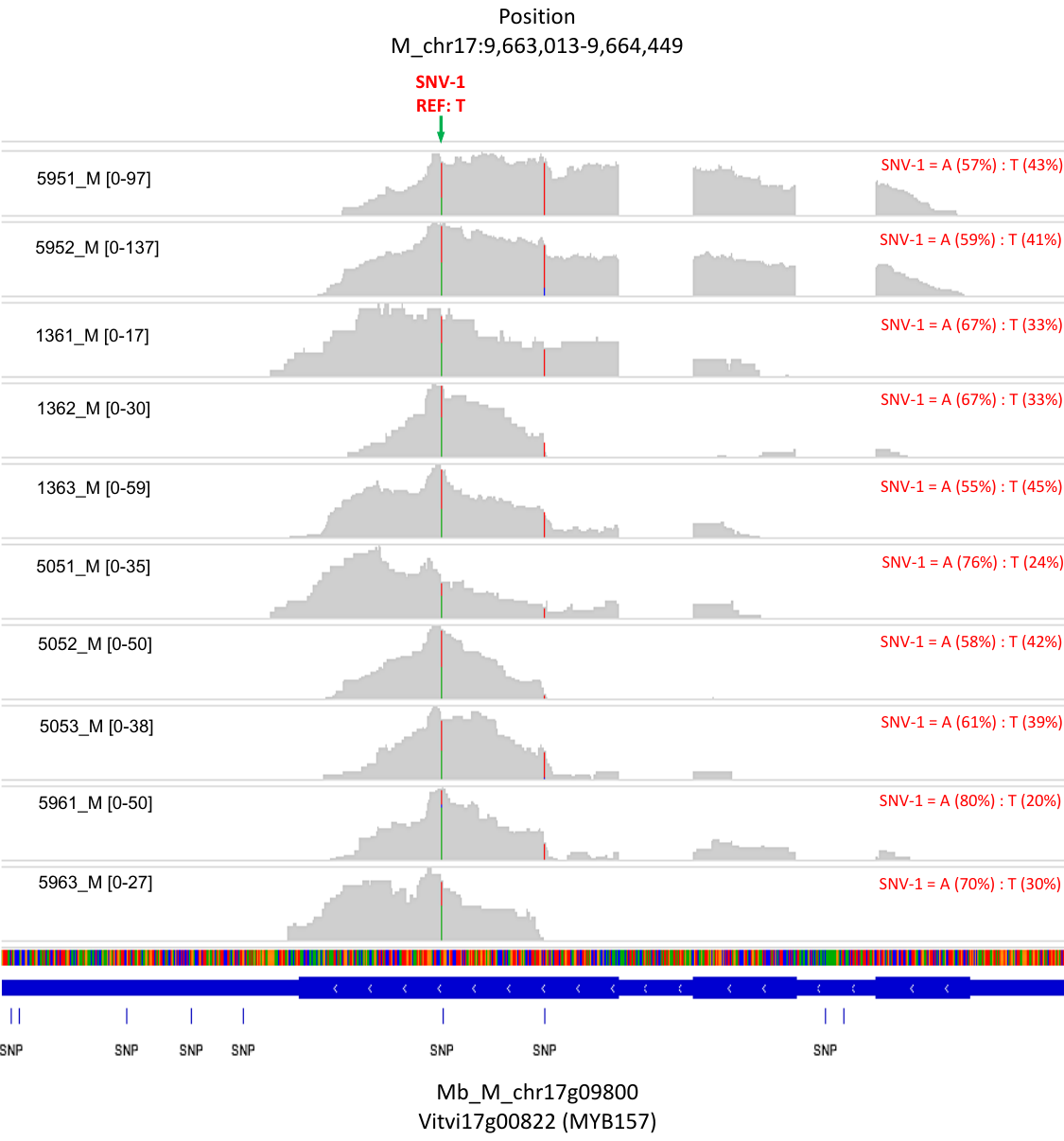

D.

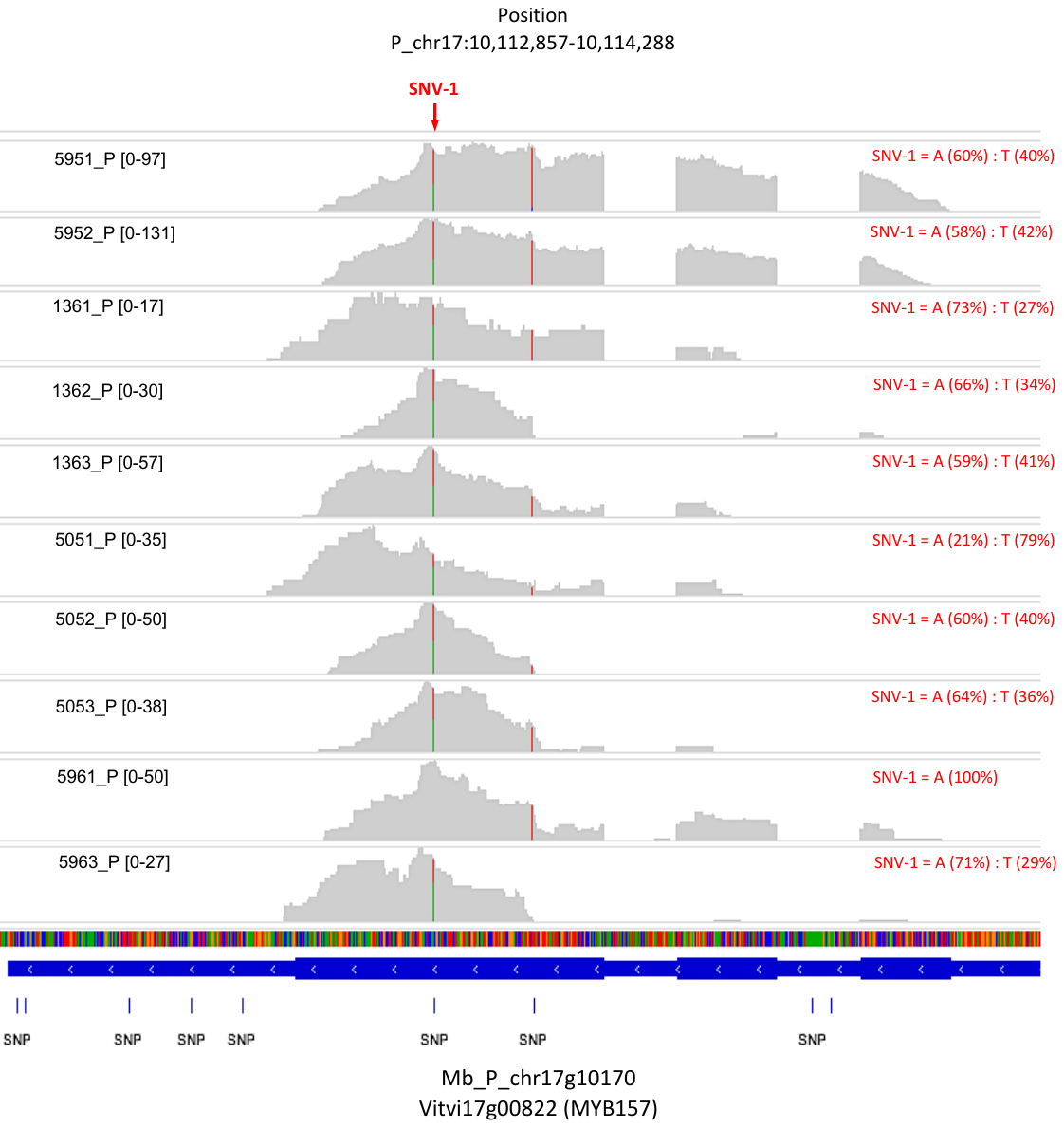

E.

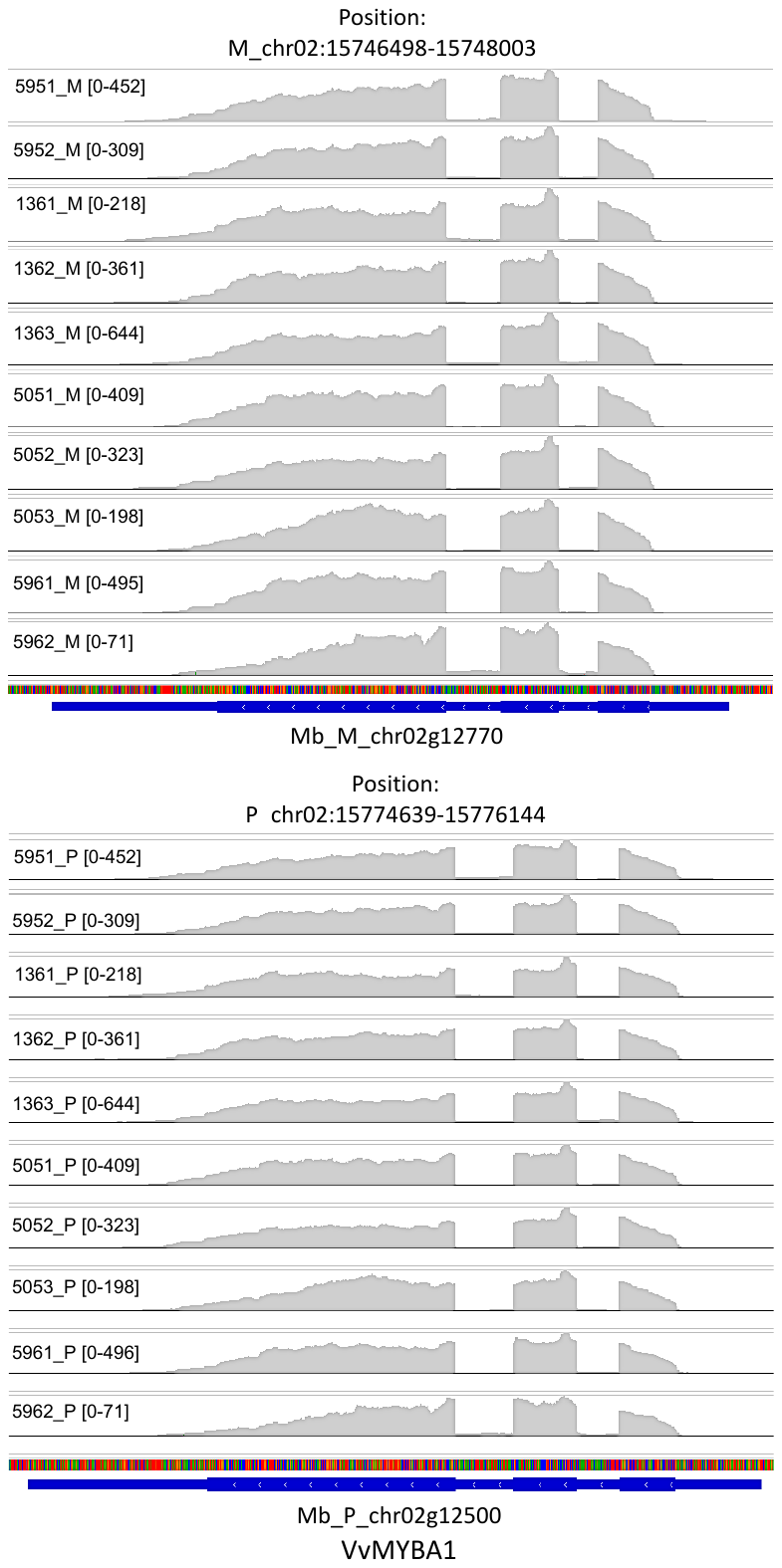

F.

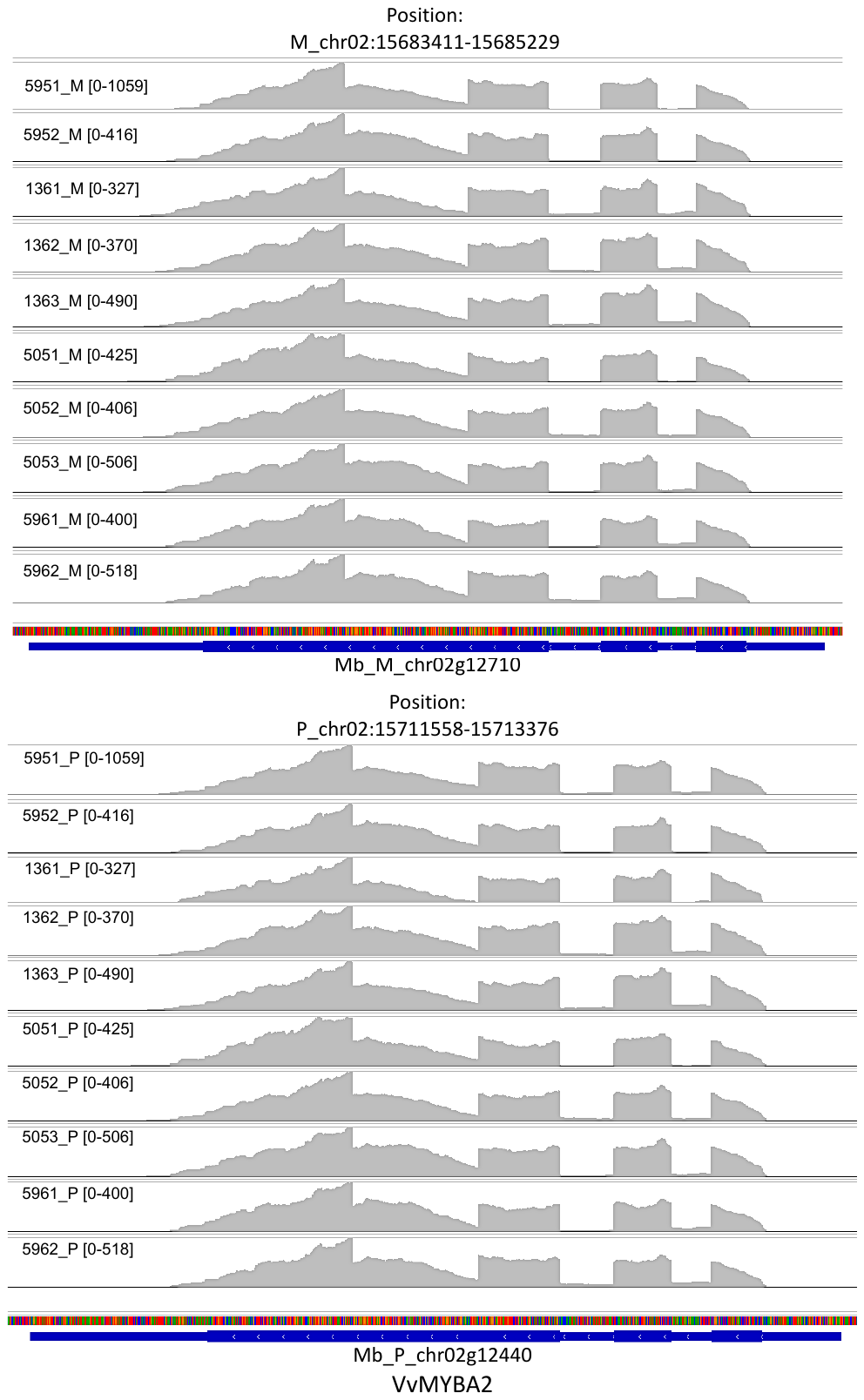

G.

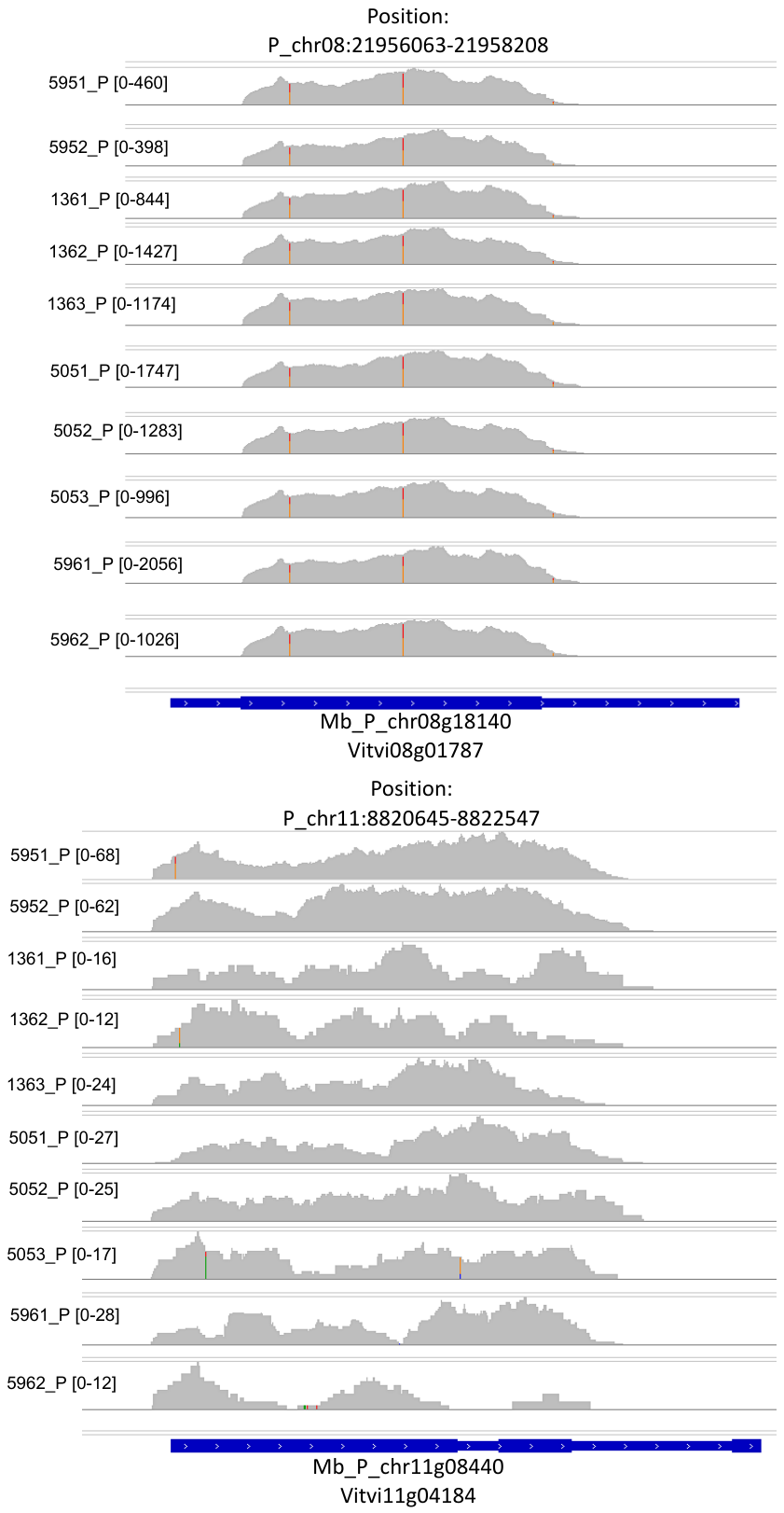
