## Supplementary material for "Diploid genome assembly of the Malbec grapevine cultivar enables haplotype-aware analysis of transcriptomic differences underlying clonal phenotypic variation": Table S1

**Supplementary Table 1**. Raw genomic data obtained to perform Malbec de novo genome assembly. **(A)** PacBio CLR reads were generated only for Malbec, from 5 SMRT cells. **(B)** Illumina PCR-free reads were obtained for Malbec and the parental cultivars. Theoretical raw coverage was estimated based on a 490 Mb genome size. **(C)** Detail of the 22 samples employed for RNA-seq experiments and for different aims in this study. Samples represent eight different clonal accessions with three biological replicates for each, except 595 and 596. In all cases RNA was extracted from whole berries (discarding the seeds) at different stages of the ripening process. Ripening berries were sampled at veraison (75% colored) and matures berries were sampled at harvest.

A.

| Cultivar | Tissue | Num. of reads | Total seqs. (Gb) | Min. length (bp) | Avg. length (bp) | Max.  length (bp) | Reads N50 (bp) | Raw  coverage |
| --- | --- | --- | --- | --- | --- | --- | --- | --- |
| Malbec | Leaves | 4,117,439 | 82.3 | 50 | 19,998 | 208,476 | 33,672 | 168x |

B.

| Cultivar | Tissue | Read length  (bp) | Number of reads (paired-end) | Total seqs. (Gb) | Raw  coverage |
| --- | --- | --- | --- | --- | --- |
| Malbec | Roots | 150 | 44,700,340 | 13.4 | 27x |
| Prunelard | Roots | 150 | 156,267,585 | 46.9 | 96x |
| Magdeleine | Roots | 150 | 158,557,983 | 47.6 | 97x |

C.

| Sample ID | Usage | Tissue | Year | Raw sequences | Raw data (Gb) | Q30  (%) | GC  (%) |
| --- | --- | --- | --- | --- | --- | --- | --- |
| 136N-1 | Annotation + Clonal diversity | Ripening Berries | 2019 | 40,848,386 | 6,1 | 93,45 | 46 |
| 136N-2 | Annotation + Clonal diversity | Ripening Berries | 2019 | 42,088,452 | 6,3 | 93,22 | 46 |
| 136N-3 | Annotation + Clonal diversity | Ripening Berries | 2019 | 47,612,222 | 7,1 | 94,28 | 46 |
| 505-1 | Annotation + Clonal diversity | Ripening Berries | 2019 | 46,591,568 | 7 | 93,53 | 46 |
| 505-2 | Annotation + Clonal diversity | Ripening Berries | 2019 | 46,541,176 | 7 | 93,68 | 46 |
| 505-3 | Annotation + Clonal diversity | Ripening Berries | 2019 | 41,308,892 | 6,2 | 92,96 | 46 |
| 595-1 | Annotation + Clonal diversity | Ripening Berries | 2019 | 50,658,656 | 7,6 | 94,31 | 46 |
| 595-2 | Annotation + Clonal diversity | Ripening Berries | 2019 | 46,924,806 | 7 | 93,08 | 46 |
| 596-1 | Annotation + Clonal diversity | Ripening Berries | 2019 | 46,340,056 | 7 | 93,86 | 46 |
| 596-2 | Annotation + Clonal diversity | Ripening Berries | 2019 | 42,838,226 | 6,4 | 92,81 | 46 |
| 53-1 | Annotation | Mature Berries | 2016 | 58,397,840 | 7,30 | 95,96 | 46 |
| 53-2 | Annotation | Mature Berries | 2016 | 56,575,042 | 7,07 | 95,91 | 46 |
| 53-3 | Annotation | Mature Berries | 2016 | 53,009,106 | 6,62 | 95,86 | 46 |
| 59-1 | Annotation | Mature Berries | 2016 | 59,256,404 | 7,40 | 95,06 | 46 |
| 59-2 | Annotation | Mature Berries | 2016 | 61,820,314 | 7,72 | 95,67 | 46 |
| 59-3 | Annotation | Mature Berries | 2016 | 48,400,140 | 6,00 | 95,95 | 46 |
| 225-1 | Annotation | Mature Berries | 2016 | 48,836,744 | 6,10 | 95,17 | 46 |
| 225-2 | Annotation | Mature Berries | 2016 | 51,107,262 | 6,40 | 95,91 | 46 |
| 225-3 | Annotation | Mature Berries | 2016 | 51,906,588 | 6,50 | 95,87 | 46 |
| 228-1 | Annotation | Mature Berries | 2016 | 48,472,008 | 6,00 | 95,67 | 46 |
| 228-2 | Annotation | Mature Berries | 2016 | 51,753,904 | 6,40 | 95,87 | 46 |
| 228-3 | Annotation | Mature Berries | 2016 | 58,565,078 | 7,30 | 95,78 | 46 |
