## Supplementary material for "Diploid genome assembly of the Malbec grapevine cultivar enables haplotype-aware analysis of transcriptomic differences underlying clonal phenotypic variation": Table S2

**Supplementary Table 2.** (A) Initial binning of Malbec PacBio CLR long-reads with Canu-Trio into Haplotypes A (inherited from Magdeleine) and Haplotype B (inherited from Prunelard), to perform the *de novo* assembly of both haplophases separately. Very few reads (0.48%) were not included in downstream analysis (too short or unassigned). (B) Summary statistics for Malbec haplophases (Malbec-Mag and Malbec-Pru) raw assemblies, after the first binning round and previous to the deduplication pipeline. BUSCO completeness refers to single copy genes and in parentheses the duplicated genes. (C) Contiguity and gene content summary statistics for the contigs that were excluded from the final assembly after deduplication pipeline. Transposable elements percentage include the types LTR, TIR, nonLTR and nonTIR, and repeat regions percentage include also centromeric and 45SrDNA sequences.

**A**

|  | Long-reads initial binning | | |
| --- | --- | --- | --- |
| Classification | Reads | Gb | Percentage % |
| Haplotype A | 1,543,308 | 35.92 | 37.48 |
| Haplotype B | 2,221,250 | 46.01 | 53.94 |
| Unknown (unassigned) | 111,995 | 0.28 | 2.72 |
| Excluded (too short) | 240,886 | 0.12 | 5.85 |
| Total reads | 4,117,439 | 82.33 | 100 |

**B**

|  | Haplophase polished contig stats | |
| --- | --- | --- |
| Metrics | Malbec-Pru | Malbec-Mag |
| Length (Mb) | 608 | 558 |
| Nº of contigs | 1,054 | 769 |
| N50 (Mb) | 6,171 | 5,628 |
| Avg. length (Mb) | 0.577 | 0.725 |
| BUSCO completeness (and duplication) (%) | 98.2 (21.7) | 98.2 (10.4) |

**C**

|  | Discarded contigs after deduplication | |
| --- | --- | --- |
| Metrics | Malbec-Pru | Malbec-Mag |
| Total length (Mb) | 123.9 | 78 |
| Nº of contigs | 940 | 616 |
| N50 (Mb) | 0.23 | 0.19 |
| Avg. length (Mb) | 0.13 | 0.12 |
| BUSCO completeness (and duplication) (%) | 23 (2.4) | 11.2 (1.1) |
| TEs (%) | 39.61 | 42.12 |
| Other repeat_regions (%) | 8.86 | 10.67 |
| Total repetitive (%) | 48.47 | 52.79 |
