## Supplementary material for "Diploid genome assembly of the Malbec grapevine cultivar enables haplotype-aware analysis of transcriptomic differences underlying clonal phenotypic variation": Table S3

**Supplementary Table 3.** Haplophases synteny evaluation and functional impact of structural variations, based on whole-genome assembly alignment (SyRi), Malbec-Pru assembly was set as reference and Malbec-Mag as the query. (A) summarizes structural and (B) sequence variation annotations. In both cases counts and cumulative lengths (in bp) for each type of variant are shown. (C) Shows a detail of how many genes overlapped with SVs between the two haplophases detected by SyRI, including copygain and copyloss variants. (D) Shows main GOs driver terms of the biological process enrichment analysis of genes overlapping with SVs (INV, TRANS, DUPs, NOTAL) using Malbec-Pru and Malbec-Mag as reference.

**A**

| Structural variations | Counts | Length reference (bp) | Length query (bp) |
| --- | --- | --- | --- |
| Syntenic regions | 6,327 | 362,146,357 | 361,963,477 |
| Inversions | 43 | 10,224,625 | 9,812,823 |
| Translocations | 2,395 | 18,865,012 | 18,558,281 |
| Duplications (reference) | 4,830 | 28,997,790 | - |
| Duplications (query) | 3,455 | - | 18,841,172 |
| Not aligned (reference) | 7,677 | 64,644,540 | - |
| Not aligned (query) | 7,594 | - | 67,238,690 |

**B**

| Sequence variations | Counts | Length reference (bp) | Length query (bp) |
| --- | --- | --- | --- |
| SNPs | 3,219,168 | 3,219,168 | 3,219,168 |
| Insertions | 324,371 | - | 3,678,071 |
| Deletions | 329,793 | 3,320,423 | - |
| Copygains | 803 | - | 2,851,269 |
| Copylosses | 802 | 3,448,980 | - |
| Highly diverged | 4,421 | 24,076,088 | 23,281,786 |
| Tandem repeats | 132 | 479,800 | 521,938 |

**C**

|  | Affected Genes | |
| --- | --- | --- |
| Structural variations | Malbec-Pru | Malbec-Mag |
| Inversions | 1864 | 1780 |
| Translocations | 1309 | 1312 |
| Duplications (reference) | 2850 | - |
| Duplications (query) | - | 2839 |
| Not aligned (reference) | 4447 | - |
| Not aligned (query) | - | 4480 |
| Copygain | 1098 | 1422 |
| Copyloss | 1896 | 1587 |
| CPG | 361 | 379 |
| CPL | 539 | 440 |

**D**

|  | source | Over-represented in SV genes term_name | term_id | adjusted_p_value |
| --- | --- | --- | --- | --- |
| Malbec-Pru as reference | GO:BP | defense response | GO:0006952 | 2,17x10^-15^ |
|  | GO:BP | DNA metabolic process | GO:0006259 | 2.15x10^-11^ |
|  | GO:BP | valine biosynthetic process | GO:0009099 | 0.02 |
|  | GO:BP | phosphorylation | GO:0016310 | 0.02 |
| Malbec-Mag as reference | GO:BP | DNA metabolic process | GO:0006259 | 1,35x10^-16^ |
|  | GO:BP | defense response | GO:0006952 | 1,88x10^-14^ |
|  | GO:BP | response to external biotic stimulus | GO:0043207 | 5,9x10^-7^ |
|  | GO:BP | glucan biosynthetic process | GO:0009250 | 0.0008 |
