## Supplementary material for "Diploid genome assembly of the Malbec grapevine cultivar enables haplotype-aware analysis of transcriptomic differences underlying clonal phenotypic variation": Table S4

**Supplementary Table 4**. Annotation and classification of repeated regions and transposable elements (TEs) across Malbec diploid genome assembly (958 Mb), performed with EDTA software. These regions were soft-masked on the assemblies for downstream gene annotation pipeline.

| TEs classes | Count | Masked (bp) | Masked % |
| --- | --- | --- | --- |
| LTR |  |  |  |
| Copia | 145,363 | 98,624,463 | 10.29% |
| Gypsy | 149,692 | 128,090,474 | 13.36% |
| unknown | 102,583 | 49,301,584 | 5.14% |
| TIR |  |  |  |
| CACTA | 72,601 | 31,965,945 | 3.34% |
| Mutator | 102,210 | 41,551,791 | 4.34% |
| PIF Harbinger | 36,685 | 14,027,356 | 1.46% |
| Tc1 Mariner | 6,696 | 1,839,269 | 0.19% |
| hAT | 26,816 | 11,950,933 | 1.25% |
| nonLTR |  |  |  |
| LINE element | 1,245 | 570,194 | 0.06% |
| unknown | 48 | 9,179 | 0.00% |
| nonTIR |  |  |  |
| Helitron | 31,804 | 11,805,467 | 1.23% |
| Repeat region | 202,989 | 74,262,677 | 7.75% |
| Totals | 878,732 | 463,999,332 | 48.41% |
