## Supplementary material for "Diploid genome assembly of the Malbec grapevine cultivar enables haplotype-aware analysis of transcriptomic differences underlying clonal phenotypic variation": Table S5

**Supplementary table 5**. (A) Pairwise comparison of proteins with orthologs among the three analyzed annotations. The analysis is based on the amino acid sequences of all the predicted proteins, including the isoforms (average of 1.2 per gene). (B) Summary statistics of the ‘per-sepcies’ orthology analysis performed with OrthoFinder comparing only both Malbec haplophases.

A.

|  | Malbec-Mag | PN40024 | Malbec-Pru |
| --- | --- | --- | --- |
| Malbec-Mag | - | 36434 | 38862 |
| PN40024 | 35887 | - | 35804 |
| Malbec-Pru | 39199 | 36737 | - |

B.

|  | Malbec-Mag | Malbec-Pru |
| --- | --- | --- |
| Number of proteins | 44,183 | 44,590 |
| Number of proteins in orthogroups | 42,397 | 42,739 |
| Number of unassigned proteins | 1,786 | 1,851 |
| Percentage of proteins in orthogroups | 96.0 | 95.8 |
| Percentage of unassigned proteins | 4.0 | 4.2 |
| Number of orthogroups containing species | 29,956 | 29,938 |
| Percentage of orthogroups containing species | 99.0 | 99.0 |
| Number of species-specific orthogroups | 317 | 299 |
| Number of proteins in species-specific orthogroups | 1,113 | 1,052 |
| Percentage of proteins in species-specific orthogroups | 2.5 | 2.4 |
