## Supplementary material for "Diploid genome assembly of the Malbec grapevine cultivar enables haplotype-aware analysis of transcriptomic differences underlying clonal phenotypic variation": Table S7

**Supplementary Table 7**. Intersection of the haplotype-specific DEGs identified in the three pairwise comparisons, for (A) Malbec-Mag and (B) Malbec-Pru. For each gene set the ortholog gene, GO id and the annotation for Biological Process (BP), Molecular Function (MF) or Cellular Component (CC) are provided if available.

A

| Intersection of DEGs detected only with Malbec-Mag haplophase | | | | | |
| --- | --- | --- | --- | --- | --- |
| Malbec-Mag | Malbec-Pru | PN40024.v4 | Change direction | GO:id | BP \| CC \| MF |
| Mb_M_chr19g01410 | Not found | Not Found | Upregulated | GO:0090502 | Protein phosphorylation |
| Mb_M_chr08g04870 | Not found | Vitvi08g04123 | Downregulated | GO:0004322 | Ferroxidase activity |
| Mb_M_chr13g10800 | Not found | Vitvi13g00981 | Upregulated | GO:0055072 | Iron ion homeostasis |
| Mb_M_chrMTg00300 | Not found | Vitvi00g04434 | Upregulated | GO:0005739 | Mitochondrion |
| Mb_M_chr19g10750 | Mb_P_chr19g11170 | Vitvi19g04350 | Upregulated | GO 0016310 | Phosphorylation |
| Mb_M_chr08g08760 | Mb_P_chr08g08960 | Vitvi08g04188 | Downregulated | GO:0046856 | Phosphatidylinositol |

B

| Intersection of DEGs detected only by Malbec-Pru haplophase | | | | | |
| --- | --- | --- | --- | --- | --- |
| Malbec-Pru | Malbec-Mag | PN40024.v4 | Change direction | GO:id | BP \| CC \| MF |
| Mb_P_chr17g01950 | Not found | Vitvi17g00152 | Upregulated | GO:000143 | Negative regulation of DNA-templated transcription |
| Mb_P_chr07g16110 | Not Found | Vitvi07g04495 | Upregulated | Not annotated | Not annotated |
| Mb_P_chr19g01400 | Not Found | Not Found | Upregulated | Not annotated | Not annotated |
| Mb_P_chr11g08440 | Not Found | Vitvi11g04184 | Upregulated | GO:0009836 | Fruit ripening \| Acyltransferase activity |
| Mb_P_chr01g04190 | Not Found | Not Found | Upregulated | GO:0007165 | Signal transduction\|  NAD+ nucleosidase activity |
| Mb_P_chr10g07750 | Mb_M_chr10g07230 | Vitvi10g00497 | Upregulated | GO:0006457 | Protein folding |
| Mb_P_chr16g02720 | Mb_M_chr16g02560 | Vitvi16g04072 | Upregulated | GO:0006520 | Cellular amino acid metabolic process |
| Mb_P_chr19g13660 | Mb_M_chr19g13010 | Not Found | Upregulated | GO:0006749 | Glutathione metabolic process |
| Mb_P_chr11g02140 | Mb_M_chr11g02180 | Vitvi11g01322 | Downregulated | GO:0006468 | Protein phosphorylation |
| Mb_P_chr08g05020 | Not Found | Vitvi08g04123 | Downregulated | GO: 0006879 | ferroxidase activity |
| Mb_P_chr08g18140 | Not Found | Vitvi08g01787 | Downregulated | GO:000762 | Phenylpropanoid metabolism |
