## Supplementary material for "Diploid genome assembly of the Malbec grapevine cultivar enables haplotype-aware analysis of transcriptomic differences underlying clonal phenotypic variation": Table S8

**Supplementary Table 8**. Detail of the core DEGs consistently detected in all pairwise comparisons, associated to the BPs enrichment that differentiated 595 from the other three clone accessions. The first four genes were enriching secondary metabolism and the other eight were enriching the response to stress GO terms. Data for the coded protein and its Molecular Function and/or Biological Process are provided, according to availability.

| DEGs - Orthology | | | Enriched GO term | | Coded protein | |
| --- | --- | --- | --- | --- | --- | --- |
| Malbec-Mag | Malbec-Pru | PN40024.v4 | Biological Process | Change  direction | Protein | Molecular Function |
| Mb_M_chr16g01980 | Mb_P_chr16g02060 | Vitvi16g00156 | Anthocyanin-containing compound biosynthetic process | Upregulated | **UDP-glycosyltransferase**(UFGT) | Anthocyanidin 3-O-glucosyltransferase |
| Mb_M_chr12g12580 | Mb_P_chr12g12840 | Vitvi12g01718 | Anthocyanin-containing compound biosynthetic process | Upregulated | UDP-glycosyltransferase  (UFGT) | UDP-glucose flavonoid 3-O-glucosyltransferase |
| Mb_M_chr13g04770 | Mb_P_chr13g04840 | Vitvi13g00350 | Regulation of Phenylpropanoid metabolic process. | Upregulated | **Ultraviolet-B receptor - UVR8** | **Chromatin-binding. UV-photoreceptor.** |
| Mb_M_chr05g19550 | Mb_P_chr05g19310 | Vitvi05g01703 | Regulation of Phenylpropanoid metabolic process. | Upregulated | **F-box/kelch-repeat protein** | **Protein binding.** |
| Mb_M_chr13g05600 | Mb_P_chr13g05620 | Vitvi13g00491 | Response to abiotic stress. Temperature heat, osmotic, salt. | Upregulated | **HSP domain-containing protein** | Protein self-association.  Protein complex oligomerization. |
| Mb_M_chr02g00360 | Mb_P_chr02g00330 | Vitvi02g00025 | Response to salt / osmotic stress | Upregulated | **HSP-like - ATPase domain-containing** | ATP-binding.  Cellular response to heat. |
| Mb_M_chr06g05190 | Mb_P_chr06g05230 | Vitvi06g01682 | Response to oxygen-containing-compound | Upregulated | **ZAT11 - zinc finger protein** | Response to chitin |
| Mb_M_chr01g05800 | Mb_P_chr01g05780 | Vitvi01g00446 | Response to salt / osmotic stress | Downregulated | **DELLA protein** | Transcription factor.  Repressor of gibberellin signaling pathway. |
| Mb_M_chr12g17200 | Mb_P_chr12g17550 | Vitvi12g02138 | Response to salt / osmotic stress | Downregulated | **HSP domain-containing protein** | Protein self-association.  Protein complex oligomerization. |
| Mb_M_chr10g18440 | Mb_P_chr10g18110 | Vitvi10g01533 | Response to oxygen-containing-compound | Downregulated | **Transcription factor SRM1** | DNA-binding.  Response to gibberellin. |
| Mb_M_chr06g12350 | Mb_P_chr06g12050 | Vitvi06g01133 | Response to oxygen-containing-compound | Downregulated | **Scarecrow-like protein 3** | DNA-binding.  Response to gibberellin. |
| Mb_M_chr01g09990 | Mb_P_chr01g10150 | Vitvi01g00845 | Response to oxygen-containing-compound | Downregulated | **Zinc finger - C2H2-type domain-containing protein** | DNA-binding.  Response to gibberellin. |
